# Classification of outcomes in antimalarial therapeutic efficacy studies with Aster

**DOI:** 10.1101/2025.04.28.651146

**Authors:** Inna Gerlovina, Sophie Berube, Jessica Briggs, Kathryn Murie, Maxwell Murphy, Amy Wesolowski, Bryan Greenhouse

## Abstract

Reliable assessment of antimalarial drug efficacy is crucial for effective response to emerging drug resistance, and therapeutic efficacy studies (TES) are the primary means of estimating *in vivo* efficacy. Accuracy of such estimates rests on correctly classifying recurrent infections developed during follow-up as recrudescences or new infections. Genotyping is used to guide classification, but polyclonal infections and alleles matching by chance make classification challenging, especially in high transmission settings. Match-counting algorithms currently recommended by World Health Organization are unreliable and produce biased results, necessitating development of principled statistical approaches. Modern genotyping methods such as multiplexed amplicon sequencing hold great potential for resolving recurrences and motivate the need for corresponding statistical methods able to utilize the rich data they provide. We propose an Adaptive Statistical framework for Therapeutic Efficacy and Recrudescence (Aster) that delivers accurate and consistent results by explicitly incorporating complexity of infection (COI), population allele frequencies, and imperfect detection of alleles in minority strains. Using an identity by descent approach, Aster accounts for alleles matching by chance and for a background infection relatedness structure that can otherwise lead to misclassification. The flexible framework can also use external information, such as parasite density and performance characteristics of a genotyping panel. Using simulations, we show that Aster dramatically outperforms match-counting algorithms in a wide variety of transmission settings and demonstrates consistently balanced performance that improves with more informative genotyping panels. Aster is implemented in a fast, fully scalable, and user-friendly R software package *asterTES* and provides accurate estimates of treatment failure for TES with any type of genotyping data, facilitating reliable evaluation of drug efficacy and effective management of malaria.

## 1 Introduction

Widespread availability of highly effective artemisinin combination therapies (ACTs) has greatly improved treatment for *Plasmodium falciparum* infections, the most fatal cause of malaria, and has contributed to a global reduction in malaria burden over the last few decades [1]. Unfortunately, progress in malaria control has recently stalled, with over 600,000 malaria deaths estimated in 2023 [2]. One of the largest threats to malaria control right now is the emergence of artemisinin partial resistance in sub-Saharan Africa, where *>* 95% of malaria cases and deaths occur [3, 4]. Treatment with a failing drug often results in partial clearance of infecting parasites, which then recrudesce,

resulting in increased morbidity, mortality, and ongoing transmission of resistant parasites. Antimalarial efficacy is primarily evaluated through therapeutic efficacy studies (TES). A TES assesses *in vivo* efficacy by enrolling patients with symptomatic malaria infections, administering directly observed drug therapy, and actively following these individuals to assess parasite clearance. Since recrudescence often occurs several weeks after therapy, extended follow-up times are required to assess for recurrent parasitemia. Importantly, the World Health Organization (WHO) recommends changing therapy if genotype-corrected failure (recrudescence) rates exceed 10%, making TES results a vital source of information to determine treatment policy [5].

In malaria endemic areas, TES participants can be reinfected during the long follow-up period. Measuring efficacy thus requires determining whether detection of asexual parasites during followup is due to an entirely new infection, which is not an indication of drug failure, or a recrudescence (possibly in combination with a new infection), suggesting true drug failure [5, 6]. In areas with high malaria transmission, including most of sub-Saharan Africa where artemisinin partial resistance is now emerging, people can be infected hundreds of times per year, and over 50% of study participants can develop new infections during follow-up [7, 8]. Results of TES in these areas – and by extension treatment policy – can therefore be dramatically affected by the accuracy of outcome classification. [9, 10, 11, 12]. The genotypes of parasites present in the blood prior to treatment (day zero [D0]) are compared with those present when recurrent parasitemia is first detected by microscopy (day of recurrence [DR]); matching parasite genotypes between D0 and DR suggest recrudescence, while the DR genotypes that are different from the ones on D0 suggest that a new infection has occurred [6]. Since *P. falciparum* populations in sub-Saharan Africa exhibit high levels of genetic diversity, it is unlikely that an individual will, by chance, be reinfected with a genetically identical parasite [6, 13, 14].

In practice, biological and technical factors complicate the comparison, compromising the accuracy of TES results [4, 15]. *P. falciparum* infections frequently contain multiple genetically distinct strains, i.e. have a complexity of infection (COI) greater than 1 [9, 16]. While genotyping can usually detect multiple alleles at each locus, it is not currently feasible to assign alleles across loci to underlying strains (i.e. the genotyping data are unphased). Since a recurrent infection would constitute a recrudescence event if it contains at least one persistent strain, regardless of the presence or absence of newly infecting strains, any matching alleles between D0 and DR at a given locus provide potential evidence of recrudescence. However, alleles from genetically distinct parasites can also match by chance, potentially resulting in misclassifying a new infection (i.e. an infection with no persistent pre-treatment strains) as a recrudescence; polyclonality exacerbates the issue by increasing the chance of matching [9, 10]. Technical issues pose additional challenges for classification: misidentification of genotyping artifacts as additional alleles (i.e. false positive alleles) can produce non-existent matches, while alleles from strains at low frequencies in the blood might go undetected (i.e. false negative alleles), potentially masking matches that are due to recrudescence [17, 18, 12].

To address these challenges, the WHO recommends genotyping 3 diverse loci and assessing allele matches between D0 and DR samples [19]. If these genotypes have at least one matching allele at all 3 loci, the recurrence is considered a recrudescence (“3/3 algorithm”) [20], otherwise it is classified as a new infection. With this algorithm, however, undetected alleles in recrudescent strains could result in misclassifying such recurrences as new infections. To allow for imperfect sensitivity of genotyping methods, the “2/3 algorithm” has been proposed as an alternative, requiring only 2 of the 3 loci to have matching alleles. These match-counting algorithms are easy to implement, but they do not explicitly account for factors that can have a large effect on the presence of matching alleles not due to recrudescence, such as allele frequency, COI, and underlying relatedness structure in the local parasite population. Resulting misclassifications lead to estimates of drug failure that may be considerably biased by factors such as transmission intensity or the genotyping method used [10, 15, 11, 4].

In light of the limitations of match-counting algorithms, the need for statistical approaches to estimate therapeutic failure rate have become apparent. Such approaches can account for biological and technical factors in a principled way, reducing bias and providing measures of uncertainty. The first likelihood-based method, CDC Bayesian algorithm [12], uses a Monte Carlo Markov chain approach and provides recurrence classification and posterior probability of recrudescence. More recently, another likelihood-based method, *PfRecur*, has been proposed [21]; it provides a posterior probability of a recrudescence event and a posterior expected proportion of recrudescent clones. Both methods are designed for genotyping data based on length polymorphisms and account for incorrectly identified and undetected alleles, assigning the same probability of detection to every allele [12] or parasite clone [21]. A different, nonparametric statistical approach [22] uses a null distribution of a genetic dissimilarity statistic between two non-recrudescent infections, approximated by empirical distribution of pairwise dissimilarity between D0 infections, to classify recurrences. This method can be used with a dissimilarity measure of the user’s preference.

Continuing development of amplicon sequencing methods, which likely have advantages over length polymorphisms in accuracy, reproducibility, accessibility, throughput, and cost [23, 24, 25, 26, 27, 28] offers potential for more reliable assessment of therapeutic failure rates. Higher numbers of genotyped loci and more reliable detection of alleles from minority strains in complex infections can improve the accuracy of estimation. Increasing generation of amplicon sequencing data motivates the need for corresponding statistically rigorous approaches that are flexible, fully scalable, able to utilize various types of information, and are easy to use. Here, we propose a novel statistical approach, Adaptive Statistical framework for Therapeutic Efficacy and Recrudescence (Aster) that performs recurrence classification along with failure rate estimation, providing inference based on an empirical Bayes approach. Aster accounts for population allele frequencies, COI, imperfect allele detection, and an underlying relatedness structure in the local parasite population (here referred to as “background relatedness”). It is further able to incorporate external information and adapt to developing genotyping methodology. The model approaches recrudescence through the concept of identity by descent (IBD), which allows us to track recrudescent and minor clones across loci. It also provides a natural way to account for the possibility of being reinfected with clones that are genetically related to the ones in a D0 sample. Aster is implemented in a software package *asterTES* [29], which is fast, numerically stable, and user-friendly, using efficient combinatorial algorithms to process unphased data. Like the framework, it is easily extendable to include new features and accomodate new inputs.

First we present the overall framework, then evaluate Aster performance against match-counting algorithms across a variety of genotyping methods and epidemiological settings using simulations. We conclude by discussing practical applications of the method.

## 2 Methods

### 2.1 Framework

Aster is designed to infer drug failure rates in antimalarial therapeutic efficacy studies. The adaptive framework consists of two main components: 1) the genetic data generating model and 2) the detection model. We assume independence between the corresponding random processes. Each of the models can be adapted to different settings and can utilize external information, allowing for flexibility in practical applications. Overlaying these models is an observation mechanism, which reflects the fact that phased genotype data are not usually directly observed; this mechanism is represented by a deterministic function. At the top level, the framework addresses recrudescence, allowing both strain| and person-specific definitions. Here we present the latter, most common approach where recrudescence is defined as an event where at least one strain from the D0 infection is recrudescent. In this case, therapeutic failure rate is the probability that an individual has a recrudescence event following therapy. First, we describe the framework components and observation mechanism for a single sample, not considering recrudescence, then expand them to a pair of samples from the same individual. The relationship between recrudescence and identity by descent (IBD) is a part of the genetic model and is described in Section 2.1.2.

#### 2.1.1 Single sample

##### Genetic data generating model

Let **X** be a *n* × *L* matrix of random variables, where *n* is COI, *L* is the number of loci, rows *X_i_**_·_***= (*X_i_*_1_*, …, X_iL_*) represent distinct parasite genotypes, and columns *X**_·_**_l_* = (*X*_1_*_l_, …, X_nl_*) represent alleles at each locus (Figure 1). **X** can be thought of as phased genotype data for a single infection. Each *X_il_* has a categorical distribution with values in a set *A_l_* = {*a_l_*_1_*, …, a_lK__l_* } of possible alleles at locus *l* and corresponding probabilities – population allele frequencies *π*(*a_lk_*). *X_il_*are assumed to be independent, which implies no linkage disequilibrium, perfect mixing of parasite strains, and no intrahost relatedness.

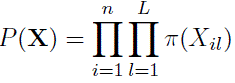

**Figure 1:**
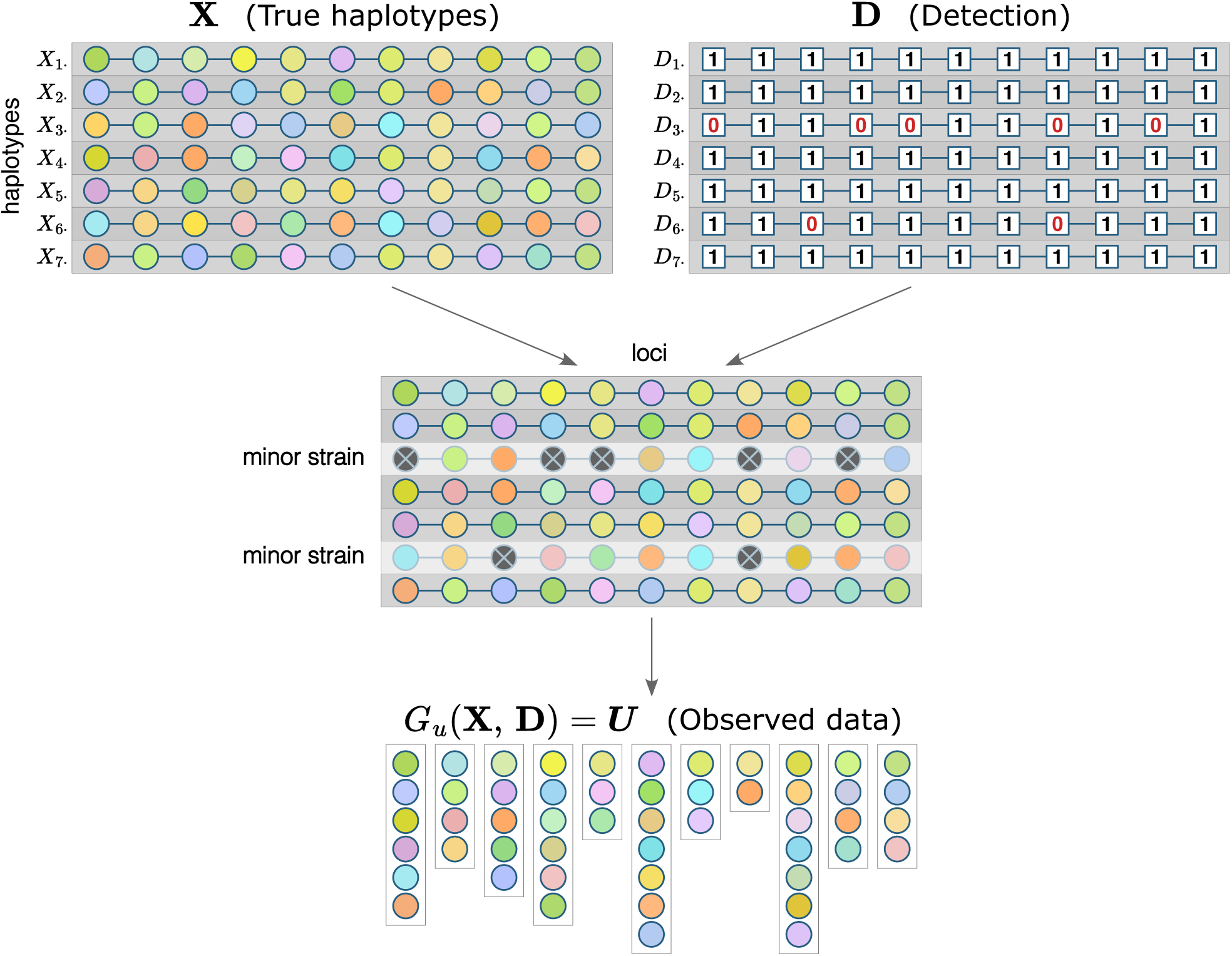
Framework components for a single sample. Each row of **X** and **D** matrices represents a strain, each column – a locus. Colors inside circles represent alleles: for each locus, the same color indicates the same allele; multiple strains can have the same allele at a given locus. Minor strains are not fully detected, which is indicated with crossed out dark gray circles, but an allele that is undetected in one strain can still be detected at a locus if it is present in another strain. Observed data for each locus are a set of detected unique alleles.

##### Detection model

In a polyclonal infection, parasite strains that are present in a sample in smaller proportions (minor strains) might be harder to detect during genotyping, and for these strains, alleles at some loci might not be detected. Let **D** be a binary *n* × *L* matrix that represents detection: if *D_il_* = 1, an allele in strain *X_i_**_·_*** at locus *l* has been detected, and if *D_il_* = 0, it has not been (Figure 1). *D_il_* are independent random variables, *D_il_*∼ *Bernoulli*(*p_il_*), where *p_il_* is a probability of detection.

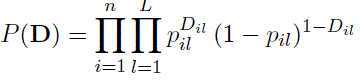

In addition to strain proportion, *p_il_* can depend on panel characteristics, overall parasite density, COI, and other factors. In many cases, it is reasonable to assume the same probability for alleles from a strain to be detected at any locus (*p_i_*_1_ = *…* = *p_iL_*), but it is possible that some markers can be more sensitive, in which case these probabilities will vary from locus to locus. Major strains can be assumed to be fully detected (*p_jl_* = 1 ∀ *l* for some major strain *j*). There might also be completely undetected strains that are not present in the sample; this special case of *p_kl_*= 0 ∀ *l* for an undetected strain *k* can be important for classification if that strain is also recrudescent. While the Aster framework formally allows such a scenario, incorporating it would require careful consideration (and possibly extending the framework) to avoid classification bias and/or extremely high uncertainty (see Supplementary Section S.4).

##### Observation mechanism

When genetic data for polyclonal infections are unphased, observed data consist of unordered lists of unique alleles detected at each locus. Let ***U*** = (*U*_1_*, …, U_L_*) be a sequence of sets that represents the observed data. ***U*** is fully determined by matrices **X** and **D**, and the observation mechanism can be represented by a deterministic function *G_u_* that maps haplotype and detection matrices to a sequence of sets of detected unique alleles (Figure 1). To define *G_u_*, we first consider a single locus. Let 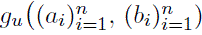) be a function that maps two sequences of the same length, the second of which is binary, to a set 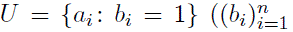 represents detection and indicates which elements of 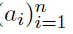 are included in *U*). Applying *g_u_* to paired columns of **X** and **D**, we get 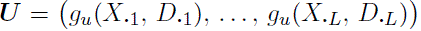, full observed data for a single sample. Thus *G_u_* is defined as 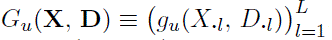. Note that if an allele from a minor strain *i* at locus *l* has not been detected (*D_il_* = 0), it can still belong to *U_l_* if it is present and has been detected in other strains.

##### Probability of observed genotyping data

Since *X_il_*⊥ *D_jm_*∀ *i, j, l, m*, the probability of observed data ***U*** for an infection with *n* distinct clones can be written simply as

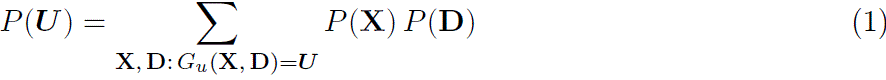

The summation in Equation (1) is taken over all the combinations of full genetic and detection matrices that are compatible with observed data, thus making this direct approach computationally unfeasible for most cases and generally unscalable; including a second sample combined with recrudescence adds another layer of complexity. While Aster uses other approaches described below to efficiently calculate the likelihood, it is a useful reference to what exactly needs to be calculated. Equation (1) does not explicitly exclude matrices with repeated rows, and there is a non-zero probability that two unrelated strains are identical by state at all the loci in a sequencing panel. However, if the panel is informative (i.e. has a sufficient number of diverse loci), this probability is small; in addition, if |*U_l_*| = *n* for at least one locus *l*, which is often the case in practice as most COI estimation methods for multiallelic data would yield estimates that are no greater than max(|*U*_1_|*, …,* |*U_L_*|), no matrices with repeated rows will be compatible with ***U***.

Using a single sample, we introduce some of the concepts of Aster implementation. First, since loci can be considered separately (while keeping in mind that for both recrudescence and detection, strains/rows need to be tracked across the loci by keeping their indices fixed), we can switch the order of the sum and the product, greatly reducing the number of combinations:

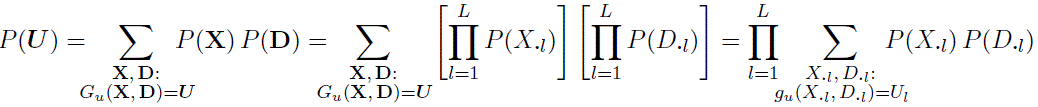

Next, we define a function that allows us to calculate the probability of a set of unique alleles in a number of strains at a given locus without explicitly allocating these alleles to specific strains. Let *P_u_*(*U, n*) be a function that calculates a probability that a sequence of length *n* has all the elements of set *U* with corresponding probabilities and only elements from set *U*: for a sequence *Z*_1_*, …, Z_n_* of independent identically distributed categorical random variables,

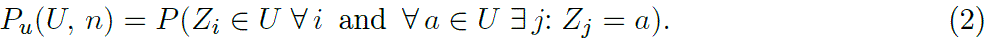

Set *P_u_*(∅, 0) = 1 (no elements in a zero-length sequence); also note that *P_u_*(*U, n*) = 0 if |*U* | *> n* and *P_u_*(∅*, n*) = 0 if *n >* 0. In Aster, elements of *U* are alleles, their corresponding probabilities are population allele frequencies, and *n* is the number of strains to which alleles in *U* belong. Supplementary section S.1 provides two efficient ways to calculate *P_u_*(*U, n*).

To incorporate missingness, recall that *P* (*D_il_* = 1) = *p_il_*. If *D_il_* = 0, i.e. an allele in strain *i* at locus *l* is not detected, *X_il_* is unknown and can be any allele *a* ∈ *A_l_*; consequently, if alleles from all the other strains j ≠ *i* are detected, *P* (*U_l_* | *n, D_il_* = 0) = *P_u_*(*U_l_, n* − 1) (see Supplementary Section S.2 for details). Then, for a sample with COI of *n* and a single minor strain *i*,

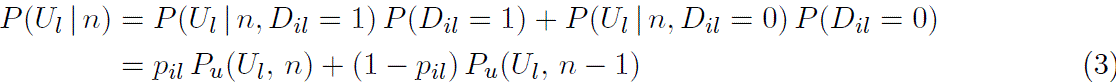

or locus *l*.

In general, for any set of minor strains,

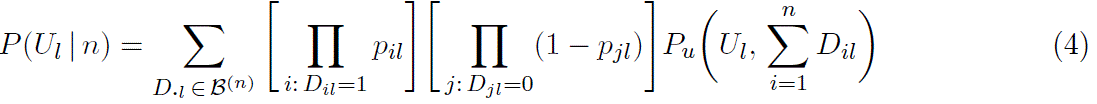

where B^(*n*)^ is a set of all the possible binary sequences of length *n* (note that the terms with *D****_·_****_l_* such that 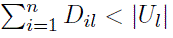 are equal to 0).

#### 2.1.2 Pair of samples and recrudescence

Recrudescence can be viewed at a person level (i.e. an individual in a study) or, alternatively, considering that many infections are polyclonal, at a strain level. Commonly, when therapeutic failure rates are estimated, a failure is defined on a person level – even if multiple strains in a single infection have recrudesced. Adopting this approach, we consider a case where at least one of the D0 strains has not completely cleared and is present in the DR sample (regardless of the presence or absence of newly accquired strains) to be a recrudescence event. Let *R_j_*be a binary random variable representing recrudescence for individual *j* where *R_j_* = 1 indicates a recrudescence event, and let *θ* be a population-level failure rate. In TES, individual responses to therapy are commonly considered to be independent, so we assume *R_j_*∼ *Bernoulli*(*θ*), *j* = 1*, …, N*, where *N* is the number of individuals in the study, to be independent and identically distributed.

##### Genetic data generating model

Let **X** and **Y** be *n_x_* × *L* and *n_y_* × *L* matrices representing D0 and DR infections from the same individual. We assume that newly infecting clones are acquired independently from recrudescence, and consequently that recurrence of parasitemia can be a result of a new infection, a recrudescence, or both. If **Y** has recrudescent strains, the rows representing them are identical to the corresponding rows in **X**, while entries *Y_jl_*in the rows of **Y** that represent newly infecting strains (if any) are drawn from the same distribution as *X_il_*. Since the order of the strains within an infection does not matter, we order them in such a way that recrudescent strains occupy the top rows of both matrices and their indices match in **X** and **Y**, e.g. if an individual has two recrudescent strains, then *X*_1_*_l_* = *Y*_1_*_l_* ∀ *l* and *X*_2_*_l_* = *Y*_2_*_l_* ∀ *l*.

We approach modeling alleles in recrudescent strains by considering identity by descent. A recrudescent strain *i* is the same in both samples, and it can be stated that *X_il_*and *Y_il_* are IBD at all loci. Let *IBD_il_*be an indicator that *X_il_* and *Y_il_*are IBD. Then, using population allele frequencies *π*(*a_lk_*),

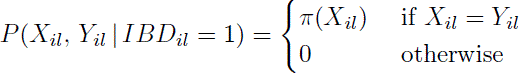

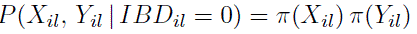

For a pair of strains *X_i_****_·_****, Y_i_****_·_***, let 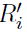 be an indicator that strain *i* is recrudescent (we want to

distinguish a strain specific variable 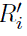 from *R*, a person-level recrudescence). If 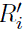 = 1, *IBD_il_* = 1 ∀ *l*; for the reverse to be true, we would need to assume that a person cannot be reinfected with strains that are related to the ones present at D0. That is not necessarily the case, and a population can have a non-zero level of background relatedness *r_bg_*, which we define as a probability that a newly infecting strain and an originally present strain are IBD at a given locus. Then

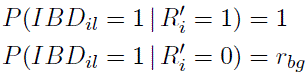

##### Detection model

Let **D***_x_* and **D***_y_* be *n_x_* × *L* and *n_y_* × *L* binary matrices representing detection for **X** and **Y**, with corresponding probabilities of detection *P* (*D_x,il_* = 1) = *p_x,il_* and *P* (*D_y,il_* = 1) = *p_y,il_*. In addition to independence between full haplotype data and detection, we assume independence between detection in two samples *D_x,il_* ⊥ *D_y,jm_* ∀ *i, j, l, m* as well as independence between detection and recrudescence/IBD status: *D_x,il_* ⊥ *IBD_il_* and *D_y,il_* ⊥ *IBD_il_*.

##### Observation mechanism

We denote observed data for samples from D0 and DR by ***U*** *_x_* = (*U_x,_*_1_*, …, U_x,L_*) = *G_u_*(**X**, **D***_x_*) and ***U*** *_y_* = (*U_y,_*_1_*, …, U_y,L_*) = *G_u_*(**Y**, **D***_y_*) respectively.

##### Likelihood

In general, for a failure rate *θ*,

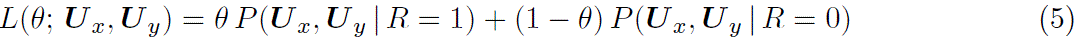

*P* (***U*** *_x_,* ***U*** *_y_* | *R* = 0) = *P* (***U*** *_x_*) *P* (***U*** *_y_*), but to derive *P* (***U*** *_x_,* ***U*** *_y_* | *R* = 1), we need to consider strainlevel recrudescence and look at probabilities with given sets of indices for minor and recrudescent strains (recall that recrudescent strain indices should match in **X** and **Y**). We start with a base case example where a possibly recrudescent strain is also a minor strain in both samples, then proceed to the general case; for derivations and special cases, see Supplementary Section S.2. Suppose there is a single strain *i* = 1 that might be recrudescent, a single minor strain *X*_1***·***_ in the D0 sample and a single minor strain *Y*_1***·***_ in the DR sample; also assume that strains *Y_k_**_·_***, *k >* 1, if any, are not related to any strains in the D0 sample. Then

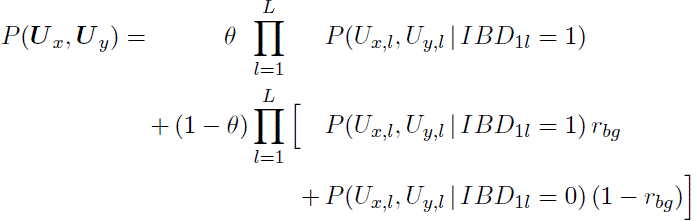

(see Supplementary Section S.3).

To find *P* (*U_x,l_, U_y,l_* | *IBD*_1*l*_), we need to account for all four combinations of *D_x,_*_1*l*_ and *D_y,_*_1*l*_ with corresponding conditional probabilities *P* (*U_x,l_, U_y,l_* | *D_x,_*_1*l*_*, D_y,_*_1*l*_*, IBD*_1*l*_):

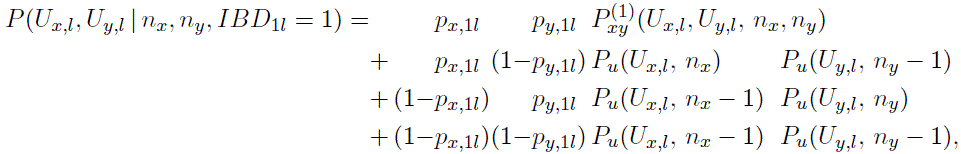

where

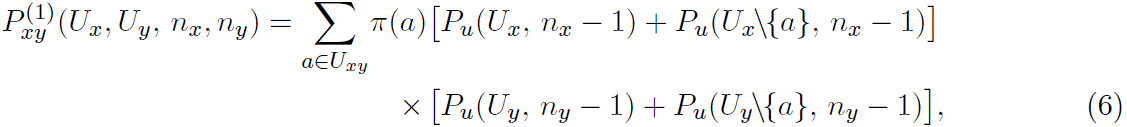

*U_xy_* = *U_x_* ∩ *U_y_*, and a set difference *U* \{*a*} is a set *U* without an element *a*. 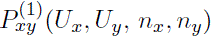 is a function that accounts for a single recrudescent strain between two samples using Equation (2). When *IBD*_1_*_l_* = 0,

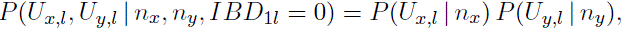

where *P* (*U_x,l_* | *n_x_*) and *P* (*U_y,l_* | *n_y_*) are calculated as in Equation (3).

In general, to accommodate any number and combination of possibly recrudescent strains, IBD pairs of strains, and minor strains, whether recrudescent or not, in both samples, we combine Equation (4) with the generalized conditional probability, which is the key component of the likelihood:

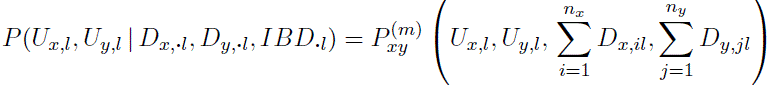

where *IBD****_·_****_l_* is a sequence 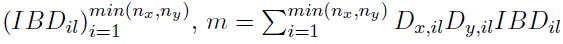, and

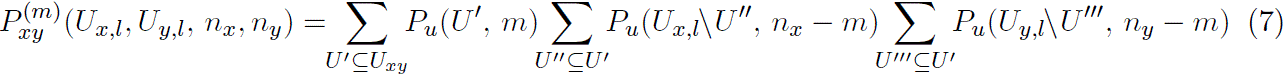

Terms with |*U* ^′^| *> m* in Equation (7) are equal to 0 since *P* (*U, n*) = 0 for |*U* | *> n*. Note that

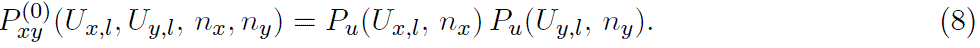

Alternatively, multisets can be used to calculate *P* (*U_x,l_, U_y,l_* | *D_x,_**_·_**_l_, D_y,_**_·_**_l_, IBD**_·_**_l_*); this approach can be more efficient in some situations. A mixed radix system algorithm [30, 31] is used for implementation of both approaches.

Equation (8) provides guidance for the treatment of missing data (no alleles detected at some loci in one or both samples). The proof that excluding such loci from the analysis does not introduce any bias can be found in Supplementary Section S.4.

#### 2.1.3 Adaptivity

The framework can incorporate different types of available information and can be combined with other methods. Depending on the amount of external information and prior knowledge, additional assumptions and constraints can be used. Finally, it can accommodate different parameters of interest and produce a corresponding output, e.g., strain-level recrudescence.

Aster explicitly accounts for population allele frequencies and COI. Allele frequencies can be estimated from D0 samples, from a combination of D0 and new infections in recurrent samples, or can be provided from larger sample estimates with information borrowed from other studies; COI can be estimated using genetic data from a single sample or a collection of samples [32]. Similarly, background relatedness can be estimated from D0 samples or its distribution can be estimated from other sources; other information (e.g., household location, travel history, or use of interventions such as bed nets) can be available to estimate the probability of being reinfected with a related strain for a particular individual. The detection model has a flexible number of parameters, where probability of detection can vary between samples, loci, and strains. They can depend on sample parasite density, COI, and characterics of the genotyping panels such as sensitivity and diversity of the markers. Prior information on therapeutic failure rates can be accommodated (and might differ for different drug combinations) with a Bayesian approach; different treatments could be compared using different priors.

Instead of a fixed value or point estimate, a distribution of *r_bg_* might be available and can be incorporated into analysis. Let *f_bg_*(*r*) be a probability density function (pdf) of *r_bg_*. Then, for the likelihood of a failure rate *θ*,

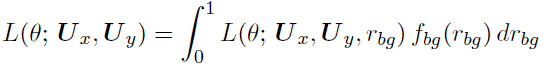

Similarly, distributions of detection probabilities can be incorporated. As an example, let the D0 sample have a single minor strain *i* with *p_x,i_*_1_ = · · · = *p_x,iL_* = *p_x_*and the DR sample have a minor strain *j* with *p_y,j_*_1_ = · · · = *p_y,jL_*= *p_y_*; let *f_x_*(*p*) and *f_y_*(*p*) be pdf’s of *p_x_* and *p_y_* respectively. Then

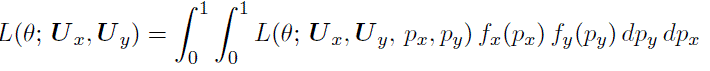

Since true recrudescence status for recurrent samples is inferred and not known for certain, there are no simple conjugate priors for the Bayesian approach. However, the posterior distribution of *θ* can be calculated numerically:

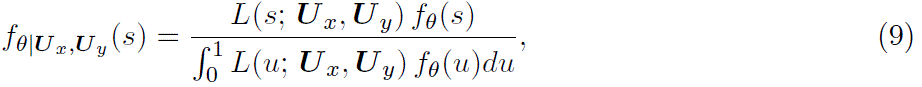

where *f_θ_*(*u*) and *f_θ_*_|***U*** *x,**U** y*_ (*u*) are prior and posterior pdf’s of *θ*.

### 2.2 Classification and inference

To classify a recurrence as a recrudescence or a new infection, we use the likelihood in Equation (5). The likelihood is linear, and classification can be based on the sign of the slope of the likelihood, or, in other words, on comparing *P* (***U*** *_x_,* ***U*** *_y_* | *R* = 1) and *P* (***U*** *_x_,* ***U*** *_y_* | *R* = 0). Formally, for a person-level recrudescence,

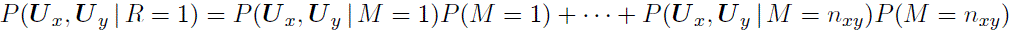

where *n_xy_* = min(*n_x_, n_y_*) and 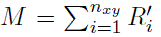 is the number of recrudescent strains. In most cases, however, it is sufficient to only consider a case with *M* = 1: if multiple strains are recrudescent while *M* = 1 is assumed, this only strengthens the evidence that *R* = 1, and if there is more evidence for *M* = 0 than for *M* = 1, it is unlikely that multiple recrudescent strains are present. While it is theoretically possible for the slopes of the likelihood functions with and without *M* = 1 assumption to have different signs (e.g. if *P* (*M* = 1) ≪ *P* (*M >* 1)), such a case is unlikely to represent a realistic scenario.

For classification inference, a useful measure would quantify the uncertainty of the individual classification, for which a posterior probability *P* (*R* = 1 | ***U*** *_x_,* ***U*** *_y_*) is a natural choice [12]. Let *A*_1_ ≡ *P* (***U*** *_x_,* ***U*** *_y_* | *R* = 1) and *A*_0_ ≡ *P* (***U*** *_x_,* ***U*** *_y_* | *R* = 0). Then

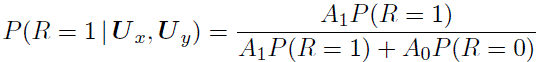

Using an empirical Bayes approach, we borrow information from all the individuals in the study by first estimating 1 − *θ* with Kaplan-Meier survival estimator 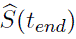 [33], where *t_end_* is the end of the follow-up period, and then using that estimate as a prior probability:

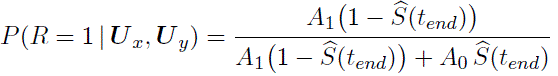

In effect, this approach provides a formal way to compare *A*_0_ and *A*_1_, weighted by prior probabilities. If we assume a prior probability *P* (*R* = 1) = 0.5, *P* (*R* = 1 | ***U*** *_x_,* ***U*** *_y_*) = *A*_1_*/*(*A*_1_ + *A*_0_). The procedure used in this approach can be iterated: recurrence classification can be updated based on the posterior probability (e.g., adopting a 0.5 probability cut-off for the classification), leading to an updated failure rate estimate. For most practical applications, a single update will be sufficient for convergence.

In addition, available external information that would contribute to prior probability or distribution can be similarly incorporated (e.g. as shown in Equation (9)).

### 2.3 Simulations

We used simulations to compare performances of Aster and match-counting algorithms for various genotyping panels, transmission intensities, genotyping error scenarios, and background relatedness levels. Panel comparisons were based on the set of 3 length polymorphisms (LP) currently recommended by the WHO (MSP-1, MSP-2, and a single microsatellite) [20], and a contemporary amplicon sequencing panel containing diverse microhaplotypes (MAD^4^HatTeR) [24]. For the amplicon sequencing panel, subsets of loci for simulations were chosen based on their heterozygosity ranking using population allele frequencies obtained from the previously analyzed datasets from high transmission areas of sub-Saharan Africa. To reflect higher genotyping error rates of LP panels, we increased missingness (1 probability of detection) twice and false positive rates 5 times compared to the amplicon sequencing panels [34, 17]. Transmission intensity levels were reflected in sample COI and the number of new infections where applicable. COI were drawn from zero-truncated Poisson (ZTP) distributions, with D0 means of 1.2, 3, and 5 for low, moderate, and high transmission settings respectively; since COI for recurrent infections are usually lower, DR means were respectively decreased by a factor of 1.8 (bounded below by 1). Genotyping errors were included with a simple “per strain” detection model for missing (false negative) alleles and a “split” model for false positive alleles, where each correctly detected allele can give rise to additional falsely detected ones ([30]).

We used three types of simulation schemes with increasing complexity: simulated pairs of samples to evaluate classification performance, simulated studies with fixed proportions of recrudescences and newly acquired infections, and studies with simulated responses to treatment combinations using an established pharmacokinetic/pharmacodynamic (PK/PD) model [35, 36, 37, 10]. With the first simulation scheme, groups of three samples were generated: a D0 sample and two DR samples, with and without recrudescence. D0-DR pairs with recrudescence were used to obtain sensitivity, while D0-DR pairs without recrudescence were used for specificity. We simulated 10,000 such triads for each combination of transmission intensity, allele detection probability, background relatedness, and genotyping panel. With the second scheme, 1000 studies were simulated for each combination. While the proportions of individuals with recrudescent strains and with newly infecting ones were fixed to remove variability due to the randomness of recrudescence events, the individuals to whom these strains were assigned were random, creating a random number of overlaps; thus the numbers of total recurrences varied. The number of recrudescent strains for each individual was drawn from a ZTP distribution with a mean of 1*/*3 of their D0 COI. For PK/PD simulations, parasite density profiles were generated using model parameters suggested in [37] with adjusted half-maximal inhibitory concentrations and maximum kill rates to achieve therapeutic failure rates at or near 0.1 [38, 39]. We used homogeneous Poisson processes to introduce new infection times with incidence per year of 6 for moderate and 12 for high transmission; the numbers of coinfecting strains were drawn from ZTP distributions with means dependent on transmission intensity (2.5 for moderate, 4 for high transmission). To mimic a TES protocol, we measured parasite density at routine time points (days 1-7, 14, 21, 28); once a detection limit of 10^8^ parasites/body had been reached, a recurrence was confirmed.

Aster analysis was conducted using *asterTES* R package [29]. Where applicable, we used naive COI estimation with a locus rank and a probabilistic COI-adjusted method ([30]) to estimate population allele frequencies for Aster. Allele detection probabilities in PK/PD simulations were estimated from sample parasite density using empirical data from mixed-strain controls for model fitting. While false positive alleles are not formally included in the Aster framework, its implementation allows for processing genetic data with false positives. When the number of alleles detected at a locus is greater than or equal to the COI, we allow for a possibility that some alleles might be false positives and therefore some might still be undetected by including a tuning parameter regulating the probability of this scenario. A small value for such probability would not affect classification results unless there is overwhelming evidence of recrudescence at other loci, in which case this evidence would be weighed against that value, and recrudescence would not be ruled out. For classification using match-counting algorithms, we calculated a proportion of loci with matches to apply these algorithms to simulated data from panels with more than 3 loci. Matches in all the loci were required for recrudescence with the 3/3 algorithm and matches in at least two thirds of the loci with the 2/3 algorithm; loci with no detected alleles in at least one of the samples were excluded from these calculations.

## 3 Results

### 3.1 Classification performance

Aster uses a probabilistic approach for recurrence classification that explicitly accounts for biological and technical factors, including COI, allele frequency, background relatedness between infections, and imperfect detection of alleles. To evaluate its performance in identifying whether a DR infection is a recrudescence or a new infection, we first simulated pairs of infections with and without recrudescences. Using different genotyping panels and a range of transmission settings, allele detection probabilities, and background relatedness levels, we compared the sensitivity and specificity of Aster to those of the 3/3 and 2/3 match-counting algorithms.

In a simulated high transmission setting, Aster demonstrated balanced performance, exceeding 0.85 specificity and 0.9 sensitivity for all simulated genotyping panels and reaching near perfect classification with an 18-microhaplotype amplicon sequencing panel (Figure 2). In contrast, the 3/3 match-counting algorithm had high specificity but much lower sensitivity, classifying many recurrences as new infections; the opposite was true for the 2/3 algorithm. Performance of match-counting algorithms worsened with increasing numbers of loci, reflecting the limiting nature of a mechanistic identity by state approach that does not benefit from additional information.

**Figure 2:**
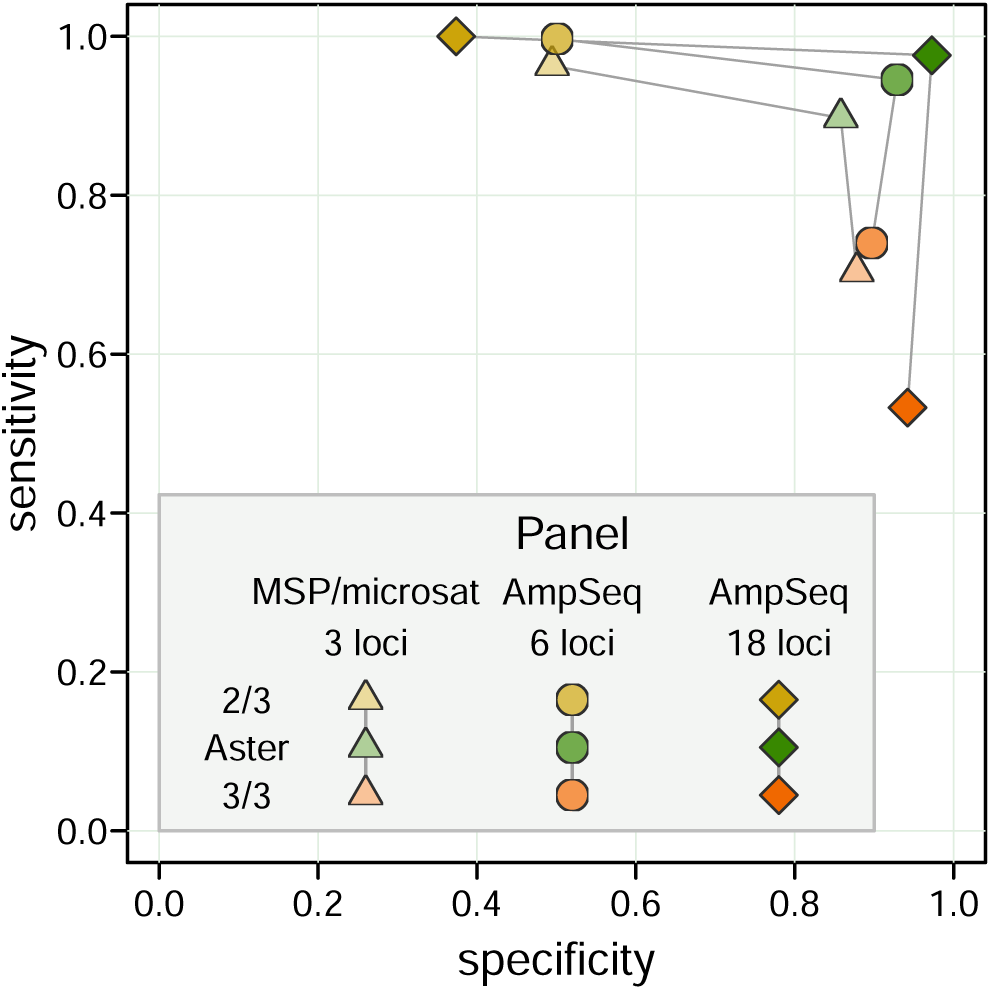
Sensitivity vs specificity for Aster, 3/3 algorithm, and 2/3 algorithm with three simulated genotyping panels: 3-locus length polymorphism (merozoite surface protein (MSP) 1, MSP-2, and a microsatellite) and 6 and 18-locus amplicon sequencing panels. A high transmission setting was used for these simulations, with a single recrudescent strain for each pair of samples and a partially detected recrudescent strain in a D0 sample; the probability of detection for that strain was set to 0.9 for all the loci in each panel. When applying match-counting algorithms to panels with *>* 3 loci, the proportion of loci with allele matches among all loci with data in both sampes was calculated; the value of 1 was required to classify a recurrence as a recrudescence by the 3/3 algorithm and of 2*/*3 or greater by the 2/3 algorithm.

Across the full range of simulations, classification was more accurate, as expected, when alleles were more likely to be detected and when transmission was lower, due to lower COI (Figure 3). However, we found that Aster had consistently higher overall and more balanced sensitivity and specificity than either match-counting algorithm. Aster performance improved with larger, more informative panels, with 6 or more loci yielding excellent accuracy in moderate transmission and 12 or more loci in most high transmission settings. Even in simulations where recurrences were the most challenging to classify, such as high transmission settings with poor allele detection, the sensitivity and specificity of Aster remained high for the 18-locus panel and improved further with 48 loci (see Figure 3). In contrast, while match-counting algorithms performed well in low transmission settings (with mostly monoclonal infections), the differences between sensitivity and specificity became pronounced at moderate and high transmission settings. This imbalance could lead to considerable overestimation or underestimation of overall failure rates, as explored below. Supplementary Figure S.1 provides similar comparisons but with varying background relatedness of infections and detection probability fixed at 0.9; as expected, higher relatedness led to decreased performance for match-counting algorithms. The performance of Aster was more robust, particularly when using larger panels.

**Figure 3:**
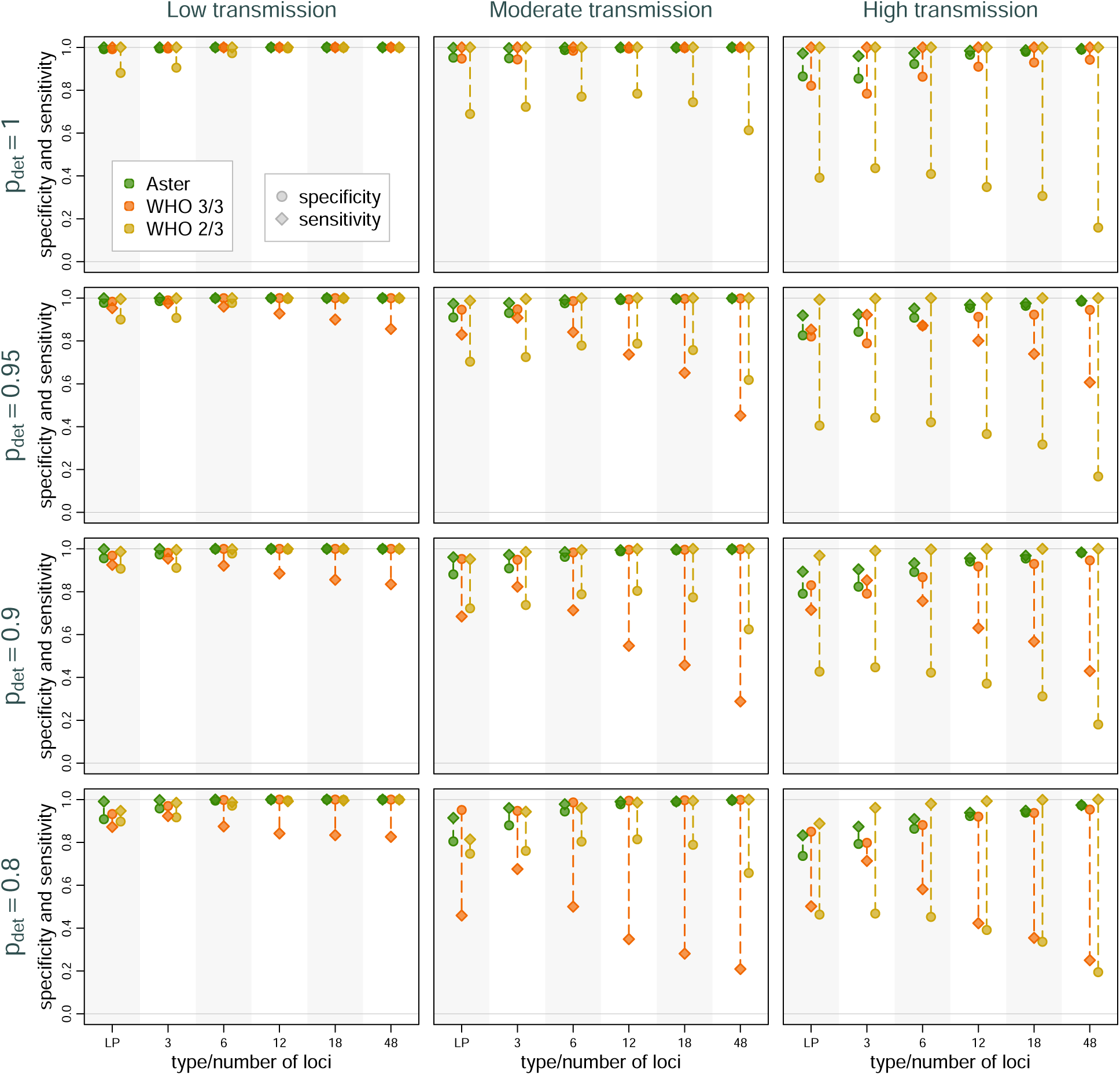
Sensitivity and specificity of Aster and match-counting algorithms across transmission intensity levels, genotyping panels, and detection probabilities (*p_det_*). The panels presented are length polymorphisms (LP, 3 loci) and MAD^4^HatTeR (varying number of loci). Dotted lines mark the difference between sensitivity and specificity for a specific method in a setting; longer lines imply greater imbalance and potentially greater bias in failure rate estimation. Background relatedness was set to 0.125.

In some cases, it may be more appropriate to include a distribution of background relatedness instead of a single value (e.g., when relatedness seems overdispersed or bimodal). As a real-world example of this type of distribution, we used the empirical distribution of pairwise relatedness obtained from D0 samples from a TES recently performed in Asayita, Ethiopia (Supplementary Figure S.2). We simulated pairs of infections with background relatedness drawn from this distribution and then classified them using Aster and match-counting algorithms. Aster provided accurate and balanced results, particularly when using more informative genotyping panels with larger numbers of diverse microhaplotype loci (Supplementary Figure S.3). In contrast, match-counting algorithms gave poor results in moderate and high transmission settings for all genotyping panels.

### 3.2 Study-level estimation accuracy

The results of a TES are interpreted at the study level, so we next simulated studies to evaluate the effect of classification accuracy on these results. First, we present the summary of the estimates obtained from studies simulated with different proportions of individuals with recrudescent and newly infecting strains. Each study had 100 individuals, a typical size for a TES. Proportions of individuals with new strains were set to 0.1, 0.3, and 0.5 for simulated settings of low, moderate, and high transmission intensity respectively. For each combination of a genotyping panel, treatment failure rate, transmission intensity level, and probability of detection, 1000 studies were simulated, and a failure rate was estimated as a proportion of individuals with recrudescences.

As expected from classification results presented earlier, failure estimates based on Aster were the most accurate and were increasingly concentrated around the true proportion of recrudescence with greater numbers of loci (Figure 4). For example, in the most challenging high transmission setting, when the true failure rate was 0.15, Aster produced results between 0.14 and 0.18 95% of the time with an 18-locus panel. In contrast, match-counting algorithm estimates were more biased, with the extent and direction of bias depending on transmission setting, true failure rate, genotyping panel, and algorithm. In general, higher transmission resulted in greater estimate variability, more bias, and greater differences between the estimates obtained with the two algorithms; notably, more informative panels led to greater bias. The 2/3 algorithm tended to overestimate failure, with dramatic overestimation in high transmission simulations, e.g. producing estimates between 0.28 and 0.41 95% of the time for a true failure rate of 0.05 when using 18 loci. The 3/3 algorithm underestimated the failure in moderate transmission, with more biased results for higher failure rates, e.g. with 95% of the estimates between 0.06 and 0.15 for a true rate of 0.2 with an 18-locus panel; the estimates were expectedly higher for high transmission, where the bias went in either direction. To evaluate the effect of imperfect allele detection on study results, we next fixed the proportion of recrudescences at 15% and considered a range of detection probabilities. Expectedly, all the methods had excellent performance when allele detection was perfect in low transmission settings. In more challenging settings, Aster results remained close to the true values (particularly for larger microhaplotype panels) for all levels of missingness, while match-counting estimates shifted downward with lower detection probabilities as more true allele matches were missed (Supplementary Figure S.4).

**Figure 4:**
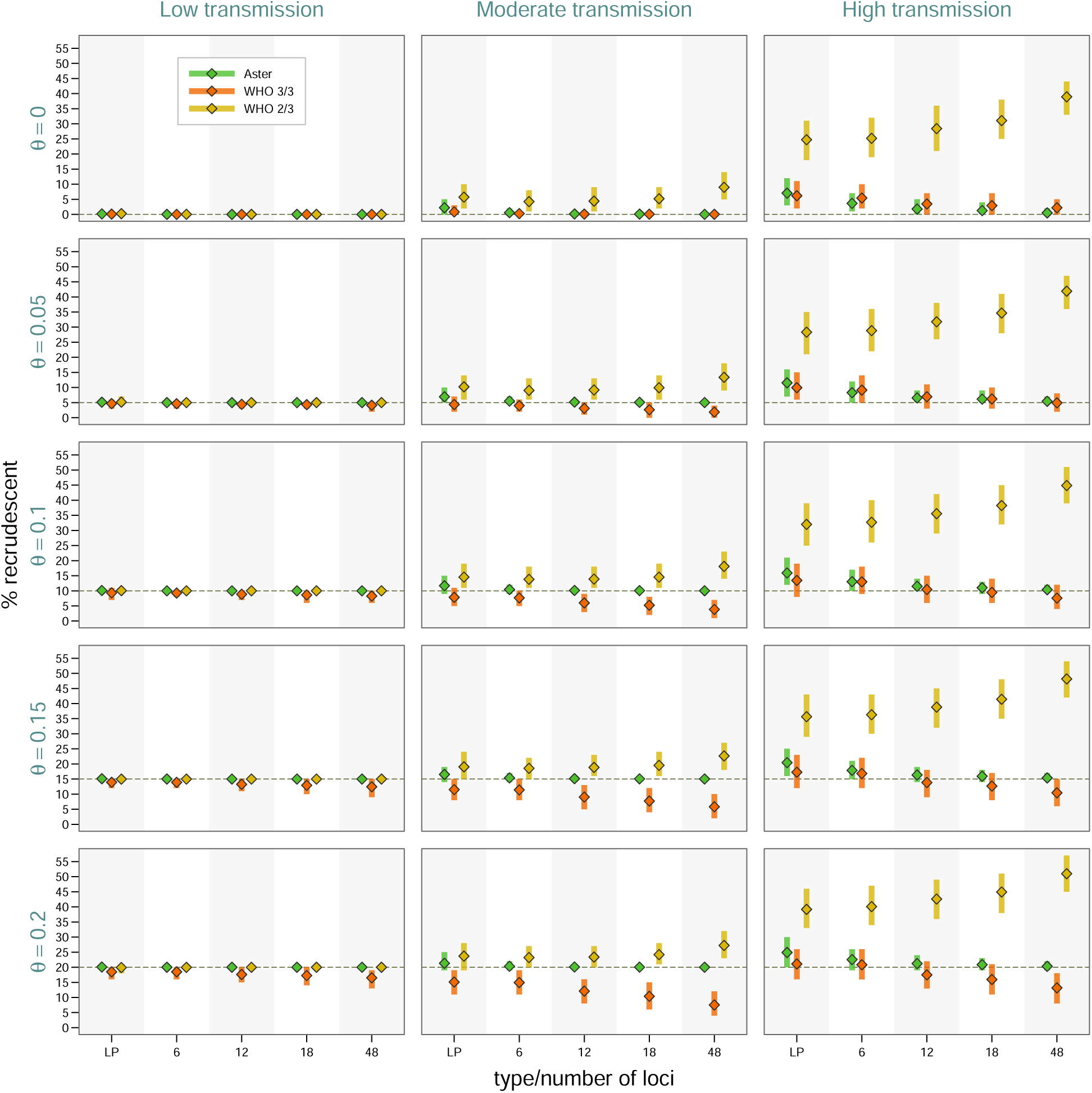
Study-level estimates across transmission intensity levels, genotyping panels, and true proportions of recrudescence events. The vertical bars represent a 95% range (0.025 to 0.975 quantiles) of the results, the diamond symbols represents a mean. Detection probability was fixed at 0.9, and background relatedness was 0.

To better approximate the underlying biological and ascertainment processes of a TES, we used a previously developed PK/PD model to simulate the dynamics of parasite clones present in D0 infections, new infections acquired during the 28 day follow-up period, and the effect of antimalarial treatment with artemether-lumefantrine (Supplementary Figure S.5). For each simulated infection where recurrence would be detected, genotyping data were generated with detection based on relative and absolute parasite density (Supplementary Figure S.6). We then applied genotyping classification algorithms to these data and calculated failure rates using the Kaplan-Meier survival estimator. The results are presented for two simulations with moderate and high transmission settings respectively, each with 1000 individuals. Comparison results were qualitatively similar to the simple simulations presented above where estimates were accurate for Aster but biased for both match-counting algorithms (Figure 5). In both simulations, the 3/3 method consistently underestimated while the 2/3 method consistently overestimated failure rates. Underestimation with the 3/3 method was more pronounced in the moderate transmission setting (producing an estimate of *<* 0.07 using 18 loci when the true failure rate was *>* 0.10), and overestimation with the 2/3 method was more pronounced in the high transmission setting (producing an estimate of *>* 0.13 using 18 loci when the true failure rate was *<* 0.10). This trend is consistent with previous results and is a likely consequence of more alleles matching by chance in the high transmission setting due to higher COI. Also consistent with previous simulation results, Aster outperforms match-counting algorithms in individual recurrence classifications (reflected in the top panels of Figure 5).

**Figure 5:**
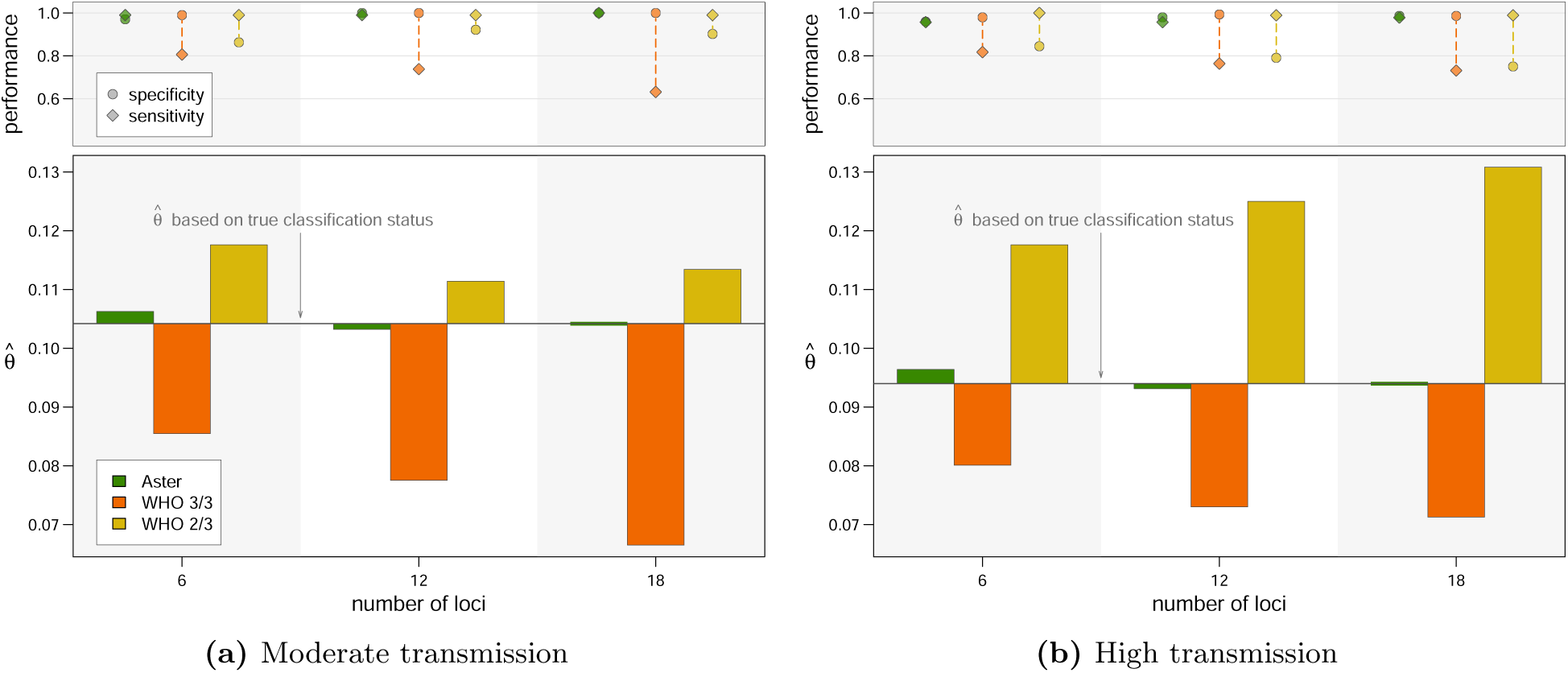
Therapeutic failure rate estimates with PK/PD model simulations using an amplicon sequencing panel with 6, 12, or 18 loci. For each figure, the bars in the lower panel represent deviation from the estimate based on true classification status; the upper panel displays performance measures. **(a)** Simulated dataset from moderate transmission: 1000 individuals with mean COI of 3.5 for D0, incidence of 6 infections/year, and an average of 2.5 coinfecting strains per new infection. **(b)** Simulated dataset from moderate transmission: 1000 individuals with mean COI of 5 for D0, incidence of 12 infections/year, and an average of 4 coinfecting strains per new infection.

### 3.3 Input estimation, parameter misspecification, and assumption violations

Since COI, population allele frequencies, detection probabilities, and other Aster model parameters are not usually known in practice and are estimated, inaccuracies of such estimates might affect Aster results. Therefore, we explored Aster’s sensitivity to misspecifications by testing the method with different scenarios. Estimating COI and allele frequencies from simulated data had a minimal effect on Aster results (Supplementary Figure S.7, rows 2, 4). Simulation of false positive alleles, though not formally included in the estimation process, had a minimal effect on amplicon sequencing results but did decrease accuracy for length polymorphisms, which were simulated with a higher false positive rate (Supplementary Figure S.7, rows 3, 4).

Sensitivity to misspecification of allele detection probability is of particular importance, since perfect detection is unlikely and missingness is generally difficult to estimate in practice. To assess the effect of misspecification, we simulated studies of 100 individuals and looked at the distribution of failure rate estimates obtained with different detection probabilities in a recrudescent strain in a D0 sample (Supplementary Figure S.8). Assuming perfect detection resulted in considerable underestimation when true detection probability was less than 1, but the effect of other misspecified values was much smaller. For example, at a true value of 0.7 allele detection (fairly poor sensitivity), assumed values between 0.8 and 0.9 still resulted in estimates of treatment failure within 0.012 and 0.018 of the true value on average; assuming perfect detection shifted the values by nearly 0.15. Thus, in all the cases when true detection was not perfect, specifying a reasonable estimate of detection probability resulted in more accurate classification than assuming no missingness at all.

## 4 Discussion

Results from TES inform national and regional antimalarial drug policies. For these results to be reliable, genotype-corrected outcomes should be consistently accurate across transmission intensities and choice of genotyping assay. We found that by explicitly accounting for population allele frequencies, COI, background relatedness, and allele detection probability within a flexible statistical framework, Aster identifies recrudescent events with high sensitivity and specificity, exceeding the performance of WHO recommended match-counting algorithms. Improvements in accuracy were particularly salient in data simulated from high transmission areas – such as those where partial artemisinin resistance is now spreading – due to higher rates of acquiring new infections and higher COI, making outcome classification more important and more challenging. In addition, as more informative data from larger genotyping panels were included, Aster estimates became more accurate – a feature that will become increasingly useful as genotyping methods continue to evolve [40, 41, 42, 43, 24].

Since Aster incorporates important factors that meaningfully impact estimates of treatment failure, we consider practical implications of connecting upstream data and external information to Aster methodology and using them as Aster inputs. The COI of infections varies with malaria transmission intensity, affecting the number of alleles matching by chance and consequently the results of traditional match-counting algorithms [9, 12, 10]. While the true COI of samples are unknown, they can be readily estimated from empirical data and incorporated into Aster to minimize bias in outcome classification. Similarly, allele frequencies will vary based on the loci genotyped and the local parasite population, affecting the probability of alleles matching. These too can be readily estimated from empirical data, for example using all or a random subset of D0 samples. Information on background relatedness between infections is explicitly included in Aster since underestimating the relatedness in a local population can lead to misclassifying a new infection as a recrudescence. As with allele frequencies, an estimate of background relatedness can be obtained from D0 samples, for example by estimating their pairwise IBD proportion, and supplied as an input to Aster. Imperfect detection of alleles may also affect results but is more difficult to estimate, potentially varying with genotyping method, COI, and parasite density. Fortunately, we show that Aster is robust to misspecification of detection probability as long as detection is not assumed to be perfect. Thus, including imperfect detection in the framework calibrates missingness in a principled way, allowing Aster to correctly classify recurrence as a recrudescence when recrudescent strains are not fully detected and would likely be missed by the 3/3 algorithm. Another type of genotyping error, false positive alleles, are not formally included in the current framework. However, while false positives can have an effect on the accuracy of match-counting algorithms, Aster is based on an IBD approach, which we demonstrate minimizes the effect of false positives on classification accuracy.

A formal framework allows Aster to approach the issue of missing data (no detected alleles at a locus, missing samples, or failed genotyping) in a systematic way. In turn, this can inform a principled treatment of right censoring and missingness in a failure rate estimation, regardless of the estimator used. For example, a recurrence with no genotyping data needs to be treated differently than a voluntary withdrawal from the study at the same time point: an individual with a recurrent infection would be more likely to have experienced a recrudescence by the end of the follow-up had they completed the study than an individual with loss of follow-up without a recurrence. In addition, the framework can be extended to include components beyond genotyping correction, for example those that deal with completely undetectable strains or persistence of gametocytes. These could affect the estimates and especially the uncertainty, more realistically reflecting the amount of unknowable information.

Reliable assessment of antimalarial efficacy is more pressing than ever, with resistance spreading in high transmission areas where accurate genotyping correction is the most challenging. Our results suggest that accuracy can be greatly improved through application of genotyping tools that provide rich genetic data, such as sensitive, diverse amplicon sequencing panels [24, 44], combined with analytical methods that take full advantage of these data and deliver consistent results across transmission levels. Fortunately, access to next generation sequencing has dramatically improved; over 80% of national public health labs in sub-Saharan Africa now have technology in place to generate high quality genotyping data (Africa CDC, personal communication, and [45]). Taking advantage of these data requires principled statistical approaches in place of simple matching-counting algorithms that can provide inconsistent results which vary by genotyping panel and epidemiological setting. Aster provides a structured statistical framework that is able to accommodate various types of information and can be amended with additional features in concordance with evolving laboratory methods and bioinformatic pipelines. The *asterTES* package, which currently provides recurrence classification with corresponding posterior probabilities and a failure rate estimate using Kaplan-Meier survival estimator, features a high level of flexibility for the user and can be extended along with the framework. With its fast and user-friendly implementation, Aster can contribute to accurate assessment of antimalarial TES, aiding stakeholders in timely determination of effective management of malaria.

## 5 Acknowledgements

We thank Ian Hastings and Sam Jones for their help and insight with PK/PD model simulations and parameter adjustments. We also thank Fitsum Girma for sharing unpublished data that provided a real life example of background relatedness from a TES. We thank Mateusz Plucinski and Andrés Aranda-Díaz for productive discussions and feedback.

This work was supported by NIH/NIAID (U01AI184646 and K24AI144048) and the Gates Foundation (INV-081860, INV-067310, and INV-048214).

## Supplementary Materials

### S.1 Calculation of *P_u_*(*U, n*)

Here we describe two approaches to calculate *P_u_*(*U, n*) that are used in implementation of Aster:

1. For *U* = {*a*_1_*, …, a_m_*} and probabilities {*π*(*a*_1_)*, …, π*(*a_m_*)}, calculate the probability of all the sequences with elements in *U*, which is 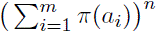, then subtract probabilities of the sequences that do not contain all of the elements in *U* using the inclusion-exclusion principal:

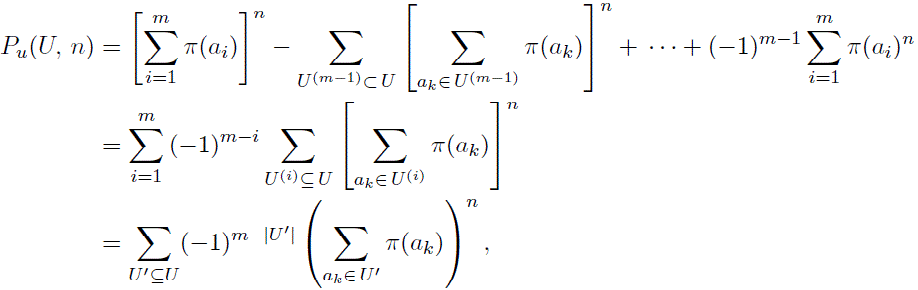

where *U* ^(*k*)^ denotes a set of cardinality *k*.
2. Generate all the multisets *S* of cardinality *n* and support *U*: |*S*| = *n, Supp*(*S*) = *U*. *P_u_*(*U, n*) is the sum of all *P* (*S*), which are calculated using multinomial probabilites.

The first approach is more computationally efficient for lower values of *m* and higher values of *n*. It gets slower with higher *m* and might encounter numerical issues (depending on a combinatorial implementation) if the orders of terms with alternating signs are much greater than the end result. The second approach is fully scalable and more efficient when *m* is close to *n*. Ultimately, the greatest efficiency is achieved with a rule prescribing which method to use in which situation, e.g. as a function of *n* and *m* that compares the number of combinations required for each method. Multisets and combinations are generated with a combinatorial mixed radix system algorithm (MIRSA) [30, 31].

*P_u_*(*U, n*) can also be defined recursively:

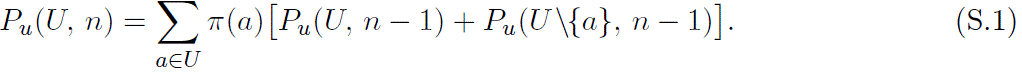

This definition follows from Equation (2) and provides an essential simplification step that will allow us to derive a conditional probability of observed data given detection and recrudescence for a pair of samples and generalize it to any combination and number of undetected alleles and recrudescent strains.

### S.2 Likelihood derivation

To translate observed unique alleles into full data and to detangle missingness and recrudescence, we need to combine combinatorics with tracking specific strains across loci (rows in **X**, **Y**) by fixing their indices. Starting with a single sample, let **D** be a random binary detection matrix such that *P* (*D_il_* = 1) = 1 ∀ *i* ≠ *k* for a fixed index *k* of a single minor strain in the sample, *p_kl_ <* 1 ∀ *l*. *D_il_, i* ≠ *k*, have a degenerate distribution and take a value of 1. *k*’th row of **D** can take values in {0, 1}. If *D_kl_*= 1 for some locus *l*, alleles in *U_l_* can be spread out across all *n* strains. If *D_kl_* = 0, those alleles have to be in the other *n* − 1 strains, and an unobserved realization of *X_kl_*could be any allele *a* ∈ *A_l_*with probability *π*(*a*). Then

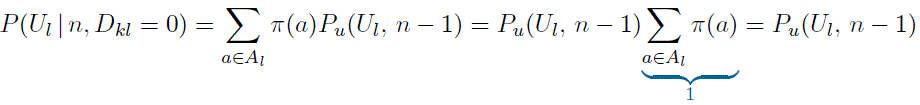

In general, for any number of minor strains, 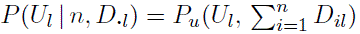.

Next, we consider a pair of samples with a single recrudescent strain *i* = 1 (the strains in **X** and **Y** are ordered in such a way that recrudescent strains are in the top rows and their indices are matching in **X** and **Y**), *IBD*_1_*_l_* = 1 ∀ *l*. For some locus *l* assume that all the alleles have been detected in both samples and that allele *a* ∈ *U_xy_, U_xy_* = *U_x_* ∩ *U_y_*is in a recrudescent strain. That means that remaining *n_x_* − 1 strains in **X** contain either all of the alleles in *U_x,l_* or all of the alleles except allele *a*; the same goes for *n_y_*− 1 strains in **Y**. Therefore, when there is no missingness,

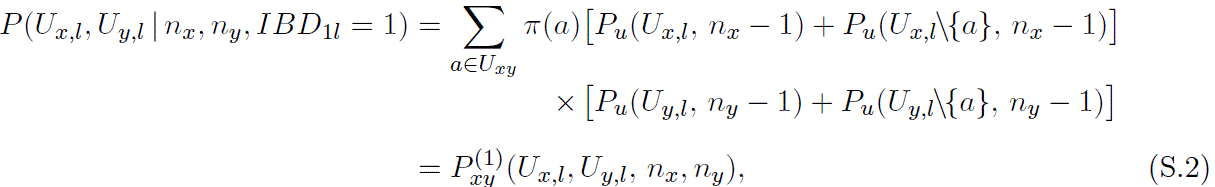

using [inlne] function defined in Equation (6). If |*U_x,l_*| = *n_x_*, an allele in a recrudescent strain cannot be in any other strains; in that case *P_u_*(*U_x,l_, n_x_* − 1) = 0. There is a similarity between Equations (S.1) (one sample) and (6) (pair of samples); Equation (6) however cannot be simplified further as the summation goes over the elements of *U_xy_* only and not over all the elements of *U_x,l_* or *U_y,l_*.

Adding missingness to this scenario, we first consider a single minor strain in each sample. At a locus, there could be no alleles missing in either sample, one sample with an undetected allele and another with detected, or both undetected. For minor strains *j* in the D0 sample and *k* in the DR sample (omitting conditioning on *n_x_*, *n_y_* in notation for brevity),

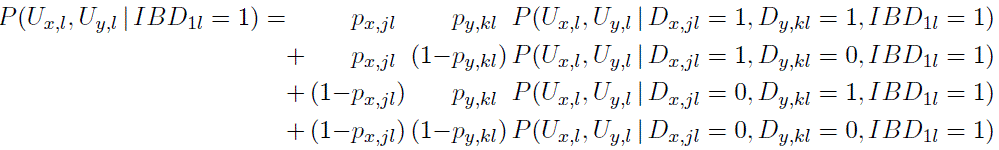

To find *P* (*U_x,l_, U_y,l_* | *D_x,jl_, D_y,kl_, IBD*_1*l*_), there are three main cases and six combinations altogether that need to be covered:

1. alleles detected in both samples;
2. allele detected in one sample, not detected in another:
  (a) an undetected allele is in a recrudescent strain;
  (b) an undetected allele is not in a recrudescent strain;
3. alleles undetected in both samples:
  (a) undetected alleles in both samples are in a recrudescent strain;
  (b) an undetected allele is in a recrudescent strain in one sample and not in a recrudescent strain in another;
  (c) undetected alleles in both samples are not in a recrudescent strain;

The first case with both alleles detected is equivalent to the one with no minor strains:

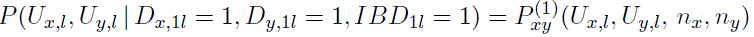

For the second case, suppose the allele in recrudescent strain *i* = 1 is missing in the DR sample. Using Equation (S.1),

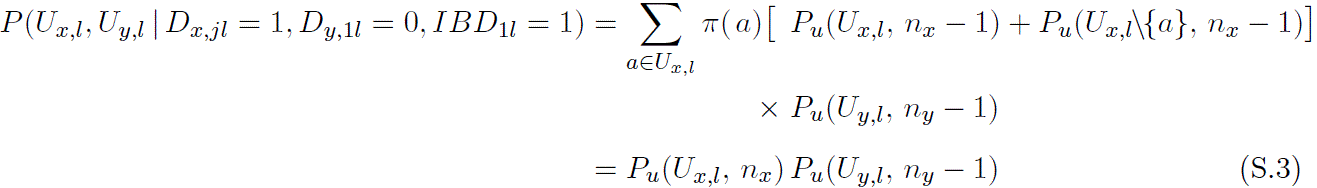

When the minor strain with an undetected allele in the DR sample is not the recrudescent one (*k* ≠ 1),

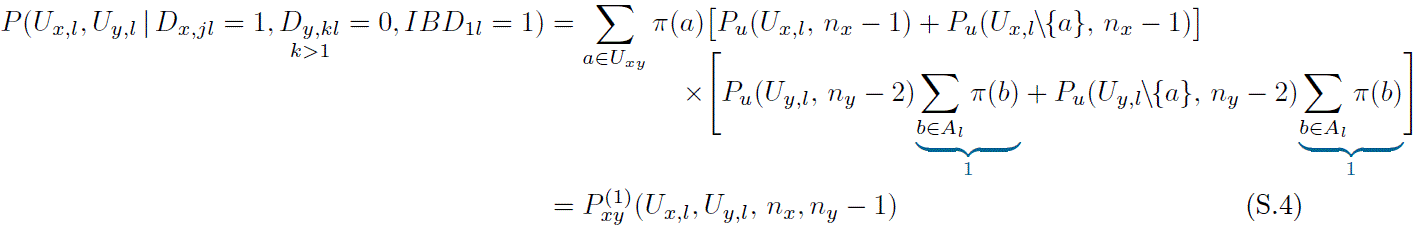

using Equation (6). Note that the index *k* is known and fixed across the loci, so when an allele *Y_kl_* is undetected, that position needs to be excluded from combinatorial calculations in 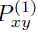 function (hence *n_y_* − 2 in the RHS of Equation (S.4)).

For the third case, when undetected alleles in both samples are in the recrudescent strain (*j* = 1, *k* = 1),

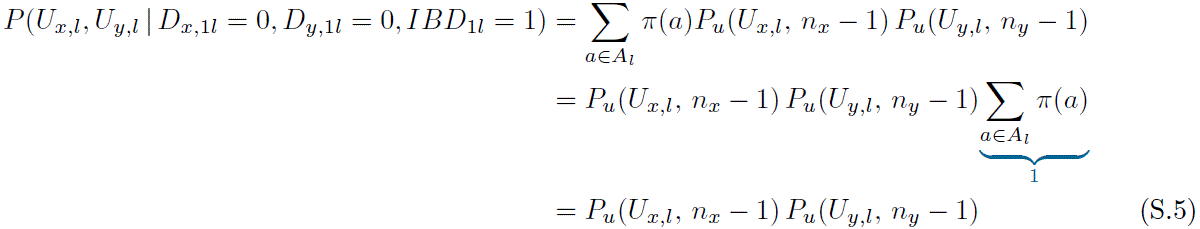

When the minor strains with undetected alleles are the recrudescent strain in the D0 sample (*j* = 1) but not the recrudescent one in the DR sample (*k* ≠ 1),

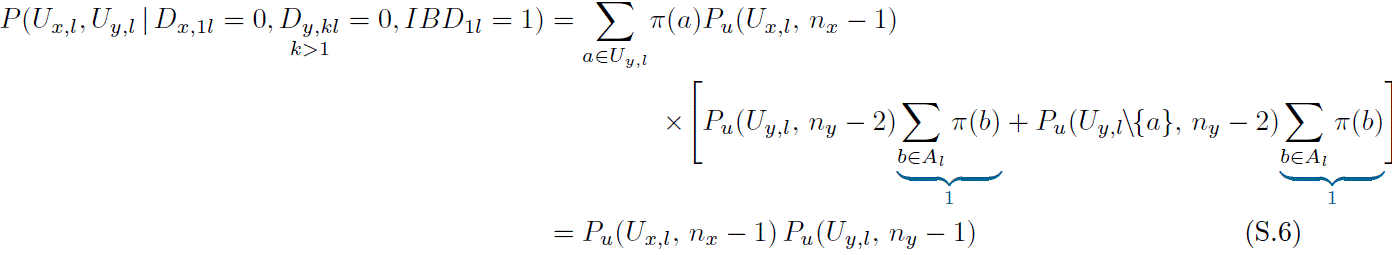

Finally, when undetected alleles are not in the recrudescent strain in both samples (*j* ≠ 1, *k* ≠ 1),

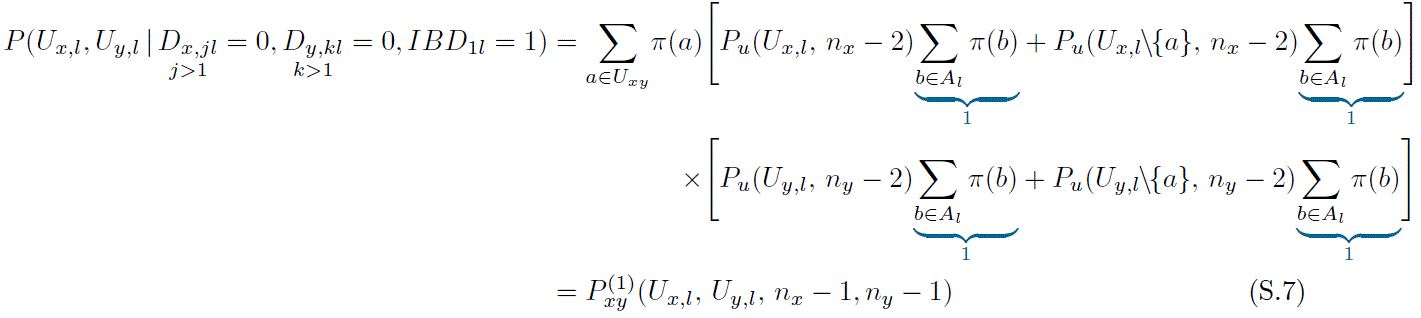

So far, we have only considered a single minor strain in each sample and a single recrudescent strain (or, more precisely, a single pair of strains that are IBD at a locus since alleles can also be IBD as a result of a reinfection with a related strain). For a general case, i.e. a conditional probability *P* (*U_x,l_, U_y,l_* | *D_x,_**_·_**_l_, D_y,_**_·_**_l_, IBD**_·_**_l_*) where *D_x,_**_·_**_l_*, *D_y,_**_·_**_l_*, and *IBD**_·_**_l_* are any binary sequences of lengths *n_x_*, *n_y_*, and min(*n_x_, n_y_*) respectively, note that the only strains that contribute to possible dependence between *U_x,l_*and *U_y,l_* are the ones where alleles are simultaneously IBD and detected in both samples; let 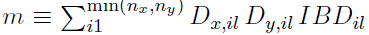 be the number of such strains. Undetected alleles, either IBD or not, factor into the conditional probability only through the second argument of *P_u_*(*U, n*) function, which makes intuitive sense (reducing the number of strains to which elements of *U* can be allocated) and is formally shown in Equations (S.3), (S.4), (S.5), (S.6), and (S.7). Therefore, the conditional probability only depends on *m* and the number of detected alleles in each sample, which can be summarized as follows:

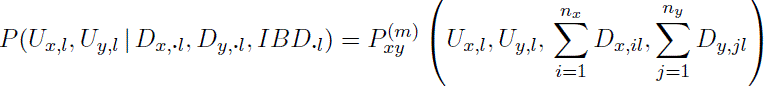

To derive 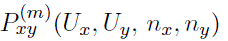, we divide strain indices in **X** and **Y** into three groups: matching indices for IBD and detected strains and remaining indices in each sample. In *m* strains of the first group, all alleles can be different, some can be the same, or all can be the same; they also do not need to include all the elements of *U_xy_*, some of which could be matching by chance. The other two groups should contain all the remaining alleles for each sample not allocated to the first group and may or may not also contain alleles already present in the first group:

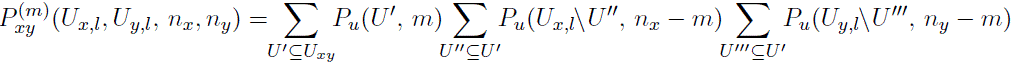

Some *U* ^′^ ⊆ *U_xy_* in the first sum can produce terms that are equal to 0: if |*U* ^′^| *> m*, *P_u_*(*U* ^′^*, m*) = 0; an empty set, which is an element of a power set of *U_xy_* will also result in a term of 0 if *m >* 0 since *P* (∅*, m >* 0) = 0. Another constraint on the cardinality of |*U* ^′^| is imposed by the cardinality of its complements with respect to *U_x,l_* and *U_y,l_*as the number of alleles not included in *U* ^′^ should not exceed *n_x_* − *m* or *n_y_* − *m*; if it does, all of the terms in the second or the third sum will be equal to 0. The cases with *U* ^′′^ = ∅ or *U* ^′′′^ = ∅ however represent scenarios where all the IBD alleles are also present in the other strains and are not generally equal to 0. Using empty sets, we obtain a special case of *m* = 0:

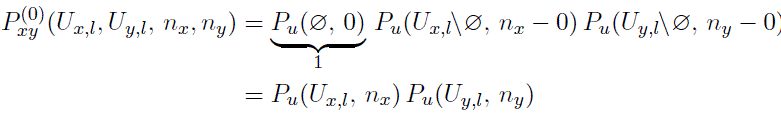

Another special case, *m* = 1, is covered in Equation (S.2) when there is no missingness; the same logic applies when alleles in some strains are undetected.

### S.3 Background relatedness

Consider a base case where at most one strain *i* = 1 can be recrudescent: 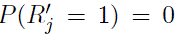 for *j* = 2*, …*, min(*n_x_, n_y_*) and consequently 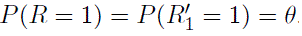. Then

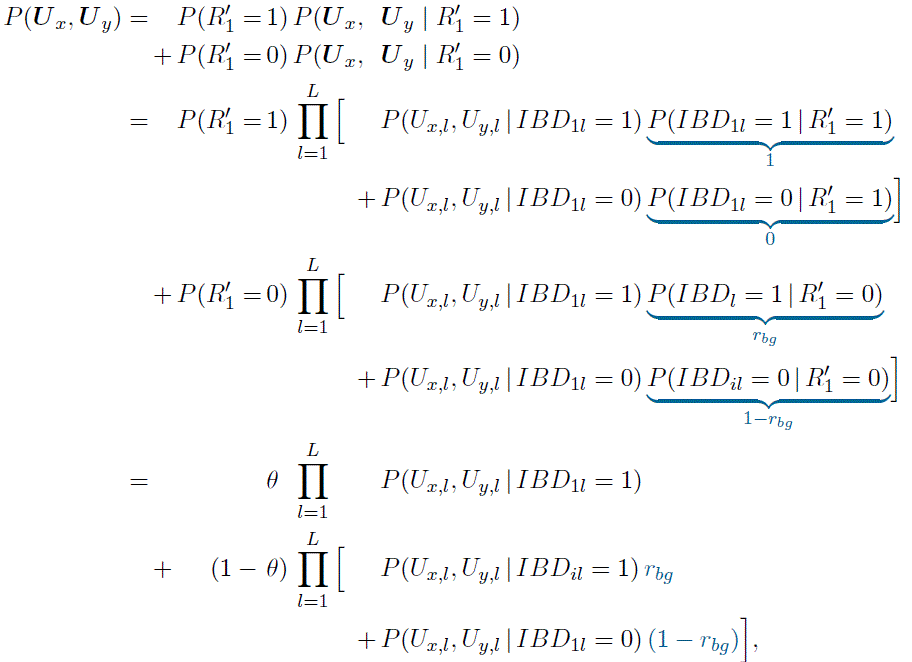

using the fact that observed data depends on recrudescence through IBD only, i.e. 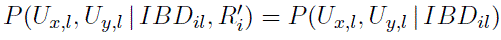.

In Section 2.1.2 we defined *r_bg_* on a strain level, which implies that every newly infecting strain can be related to an originally present strain with *r_bg_*IBD on average. If multiple pairs of nonrecrudescent strains are related between a pair of the D0 and DR samples, all the IBD combinations at a locus need to be accounted for. In addition, an assumption of independence of all interhost *IBD* variables at a locus would be required to avoid transitive dependencies. In practice, however, relatedness for a pair of samples is often estimated on an infection level. This interpretation can be easily accommodated in Aster by restricting interhost relatedness to a single pair of strains, which would allow infection-level estimates to be used for *r_bg_* directly while decreasing the number of combinations required to calculate the likelihood.

### S.4 Missing data

Missing data *U_x,l_*= ∅ or *U_y,l_*= ∅, i.e. no detected alleles at locus *l* in a sample, is a special case of realization of *D_x,_**_·_**_l_* or *D_y,_**_·_**_l_* that is actually observed (a sequence of 0’s); another special observed case being a sequence of 1’s, which would be inferred when |*U_x,l_*| = *n_x_* (assuming no false positive alleles). Intuitively, such locus provides no information as to recrudescence regardless of detected alleles in the other sample; we show this formally. Since 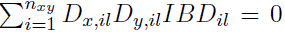 regardless of 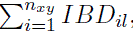, where *n_xy_* = min(*n_x_, n_y_*), we use Equation (8) and *P*(∅, 0) = 1 to get

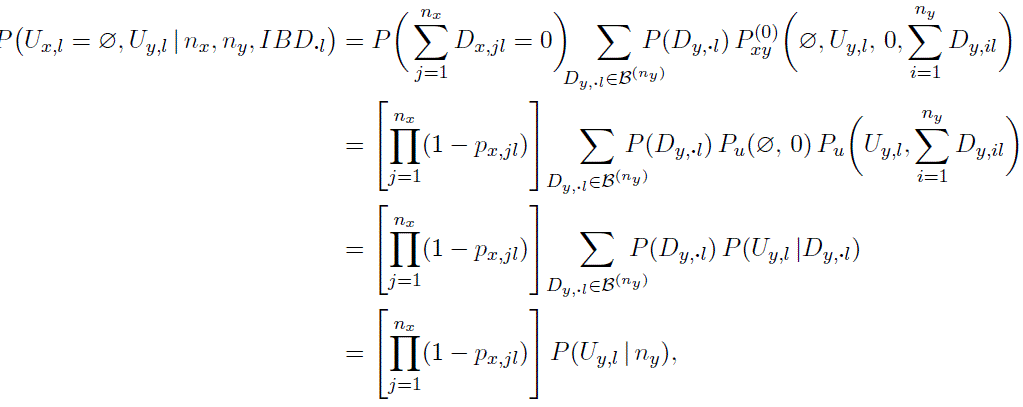

where *B*^(^*^n^*^)^ is a set of all the binary sequences of length *n*. Similarly,

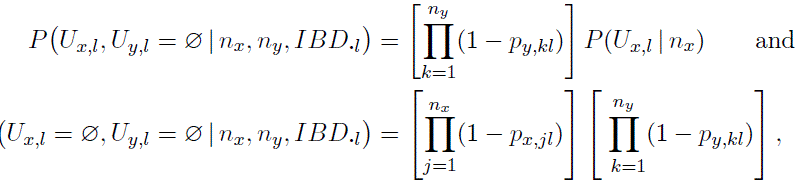

the latter being a special case when no alleles are detected at locus *l* in either sample. The fact that 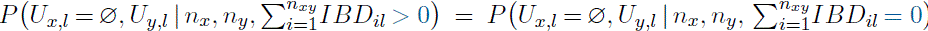 means that recrudescence and no recrudescence are given the same weight, which can be factored out of the likelihood (of the form *θ C*_1_ + (1 − *θ*) *C*_0_) and thus does not affect the shape of the log-likelihood function. This confirms that loci with no data in at least one of the samples can be ignored without introducing any bias.

Using similar logic, consider another special case that warrants attention and might be potentially problematic – completely undetected strains if they are recrudescent. Let strain *i* = 1 be possibly recrudescent and let *p_x,_*_1_*_l_* = 0 ∀ *l*; assume no other strains can be recrudescent or IBD. Then 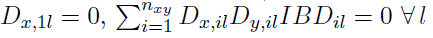, and

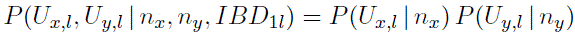

regardless of the value of *IBD*_1_*_l_*. Since that is the case for all the loci, *P* (***U*** *_x_,* ***U*** *_y_* | *R* = 1) = *P* (***U*** *_x_,* ***U*** *_y_* | *R* = 0), which implies that recurrence classification cannot be performed with any degree of certainty, and the individual essentially does not contribute to failure rate estimation (apart from censoring). This situation presents a potential source of bias: incorrectly assuming no recrudescent strains to be completely missing might lead to underestimation of drug failure, and always assuming that there is an undetected recrudescent strain will never allow a recurrence to be classified as a new infection. Fortunately, this is not likely to occur commonly; if available, an estimate of the probability of this occuring could be incorporated into the framework.

### S.5 Supplementary Figures

**Figure S.1:**
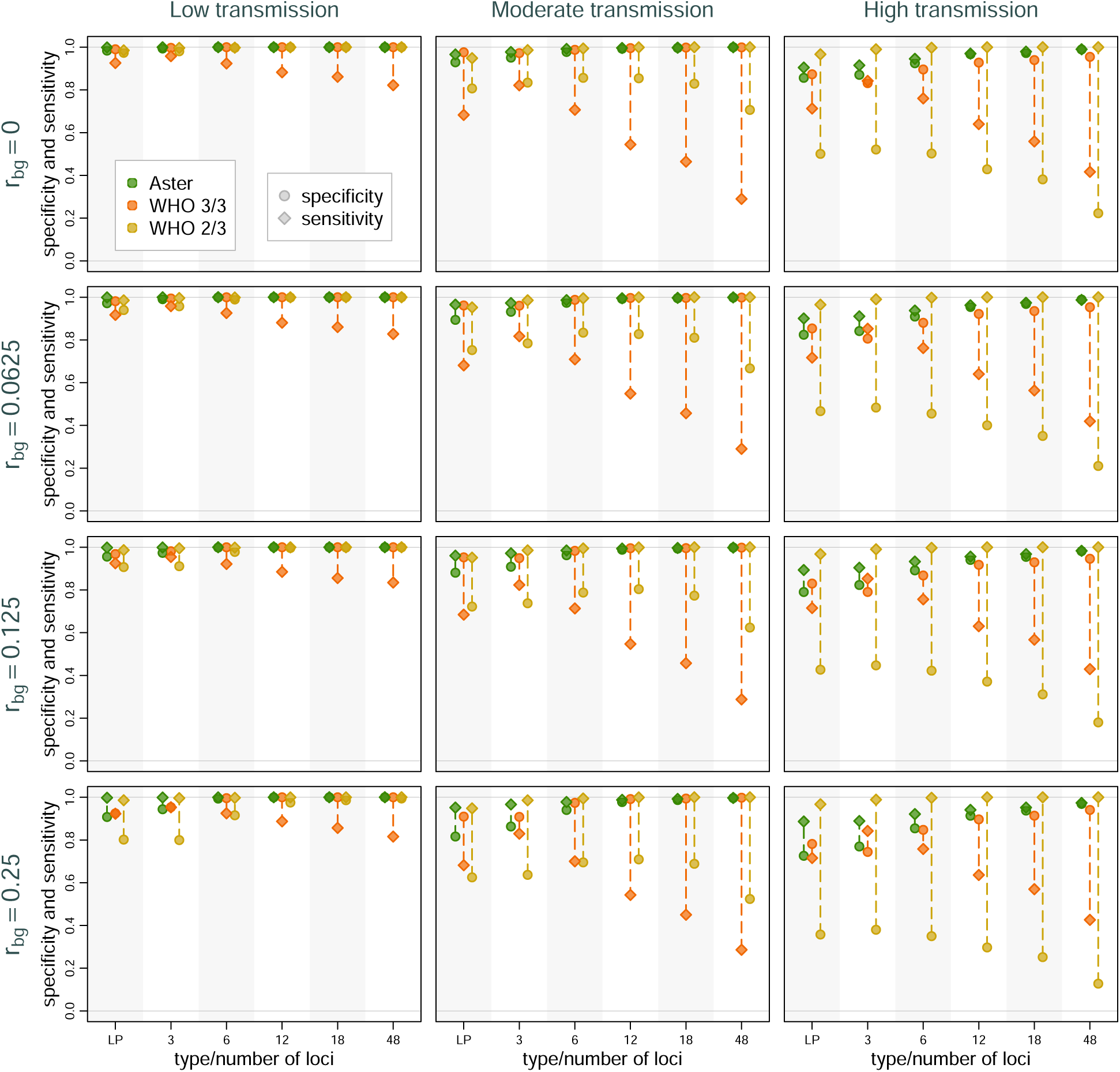
Sensitivity and specificity of Aster and match-counting algorithms across trasmission intensity levels, genotyping panels, and background relatedness (*r_bg_*) levels. Dotted lines mark the difference between sensitivity and specificity for a specific method in a setting; longer lines imply greater imbalance and potentially greater bias in failure rate estimation. Detection probability was set to 0.9.

**Figure S.2:**
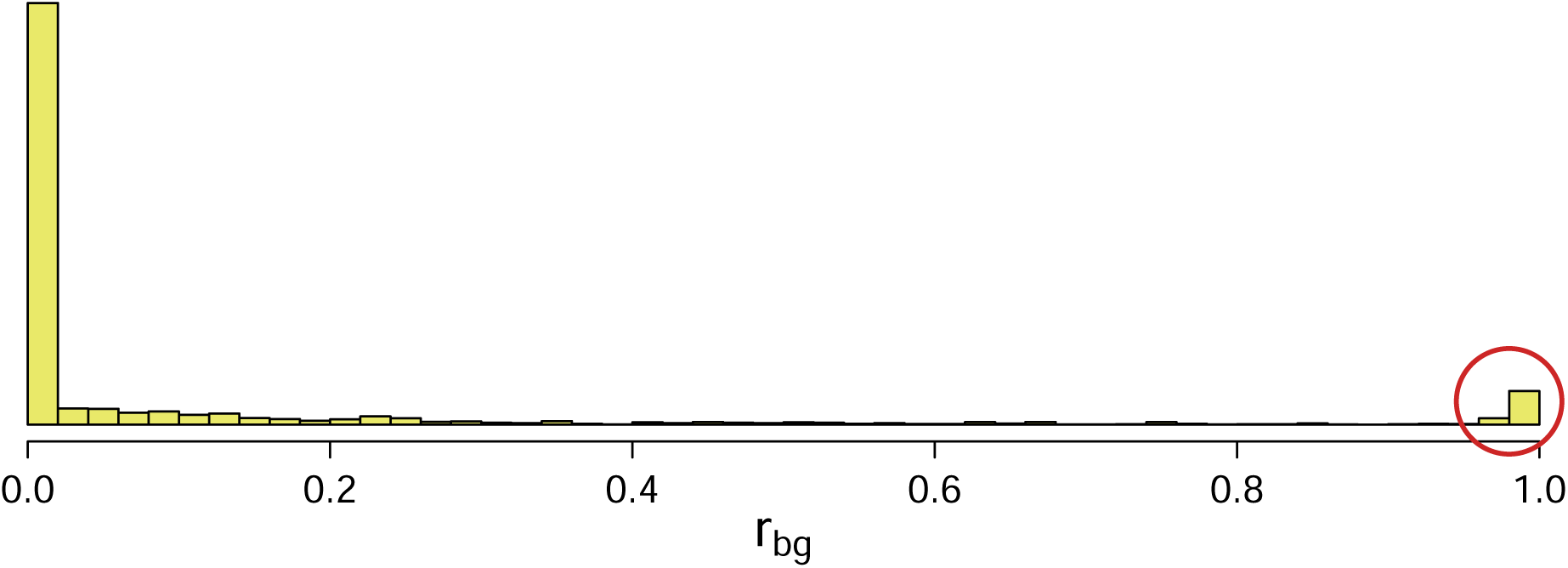
Histogram of pairwise sample relatedness in Asayita, Ethiopia, estimated using Dcifer [30]. A red circle highlights an unusually high proportion of sample pairs that were highly related.

**Figure S.3:**
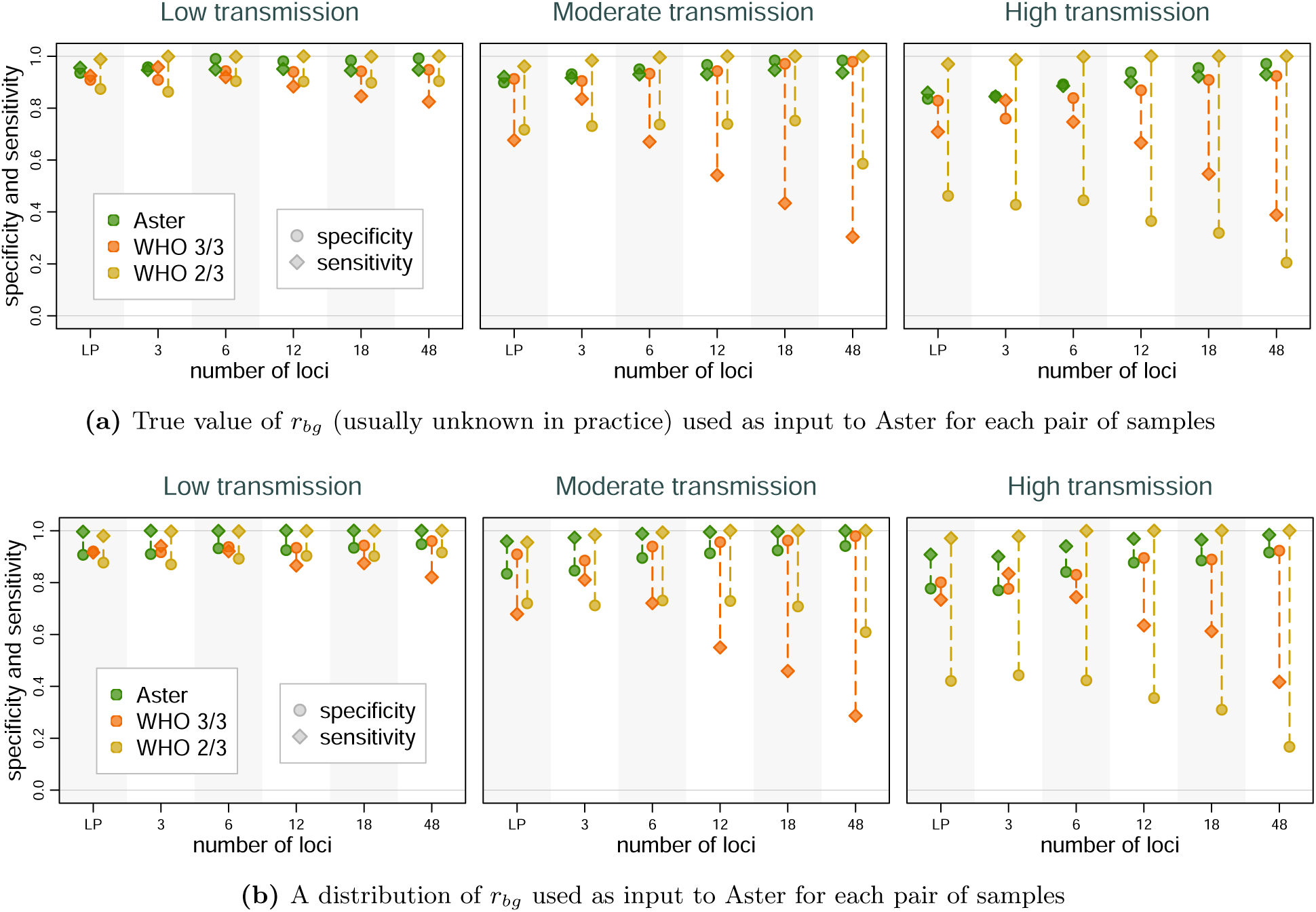
Classification performance of Aster and match-counting algorithms across genotyping panels and transmission intensities from simulations using an empirical distribution of *r_bg_*from Asayita, Ethiopia. Detection probability was set to 0.9.

**Figure S.4:**
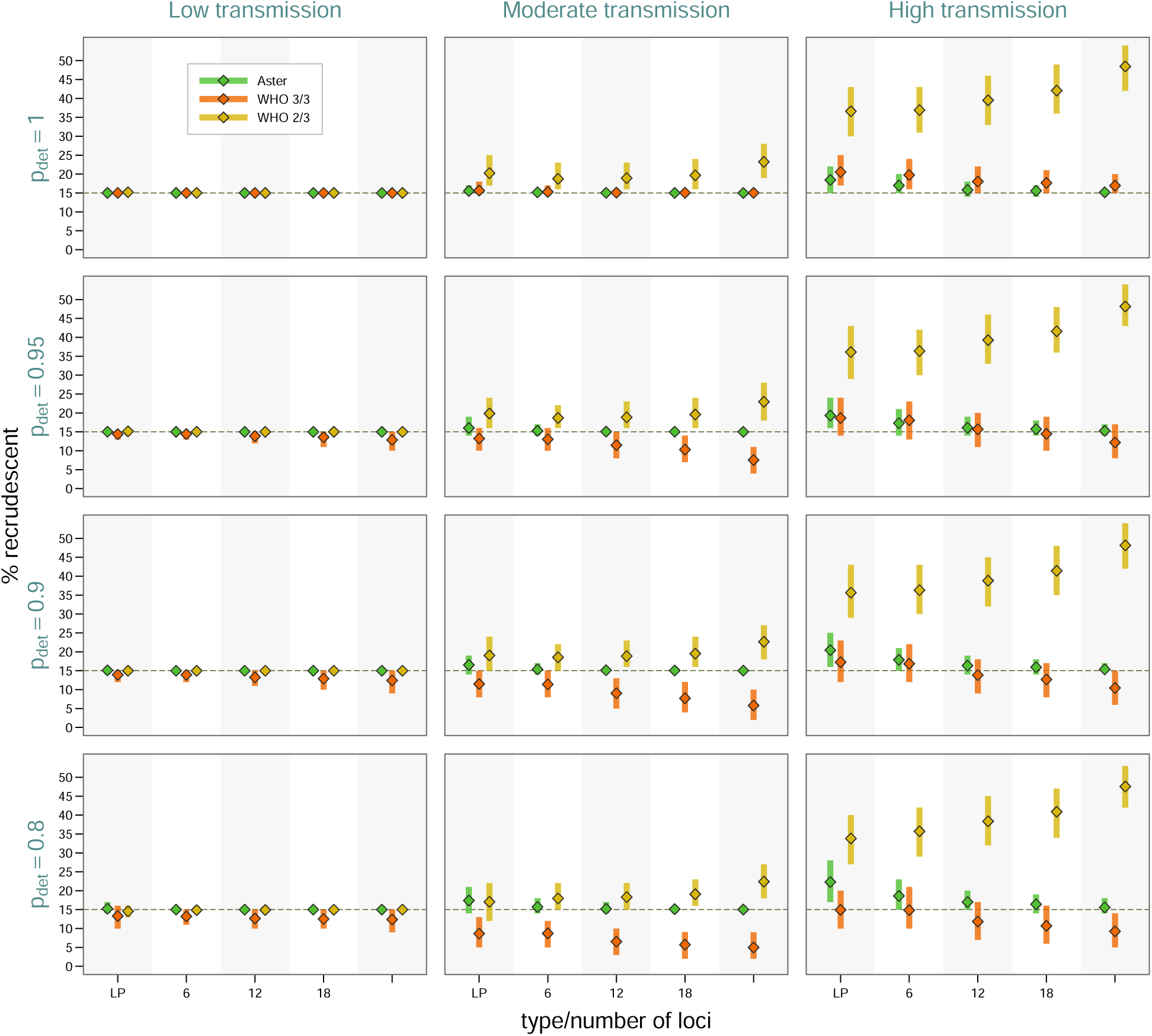
Study-level estimates across transmission intensity levels, genotyping panels, and detection probabilities (*p_det_*). The vertical bars represent a 95% range (0.025 to 0.975 quantiles) of the results, the diamond symbols represents a mean. Proportion of recrudescent events was fixed at 0.15 and background relatedness was 0.

**Figure S.5:**
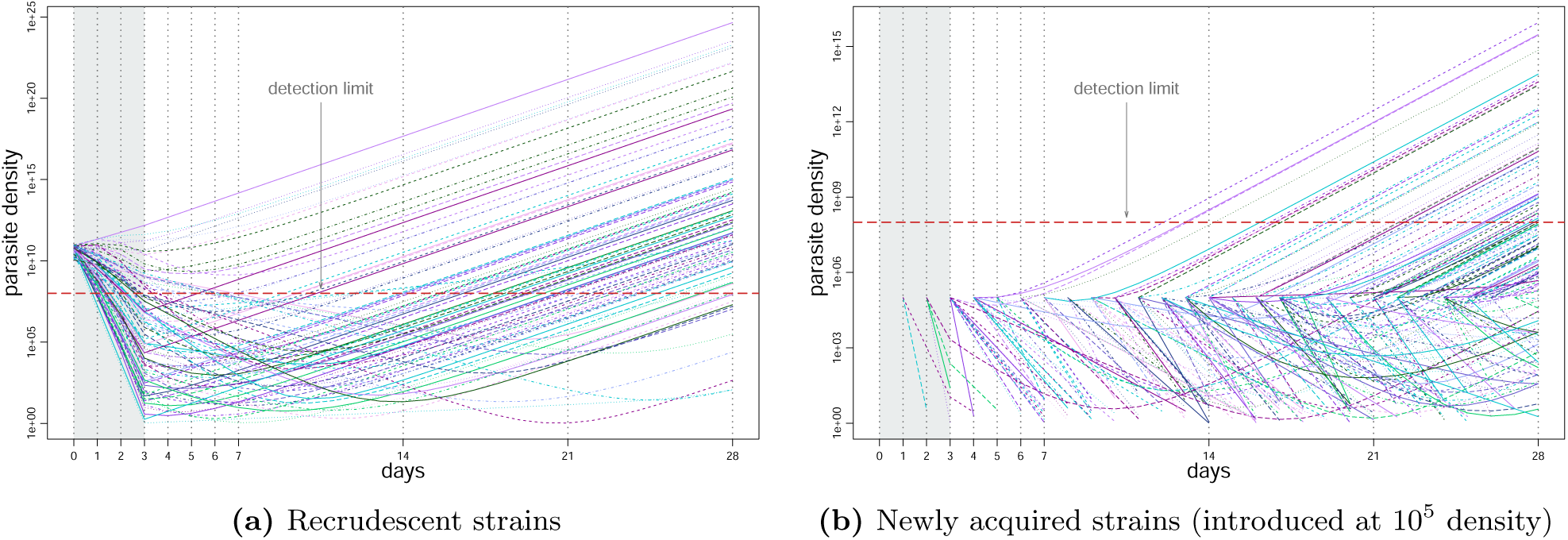
Simulated per-strain parasite density dynamics from PK/PD model. Vertical dotted lines represent time points at which the sample parasite density is measured; shaded area represents a treatment period. A dotted red horizontal line indicates a per-sample detection limit.

**Figure S.6:**
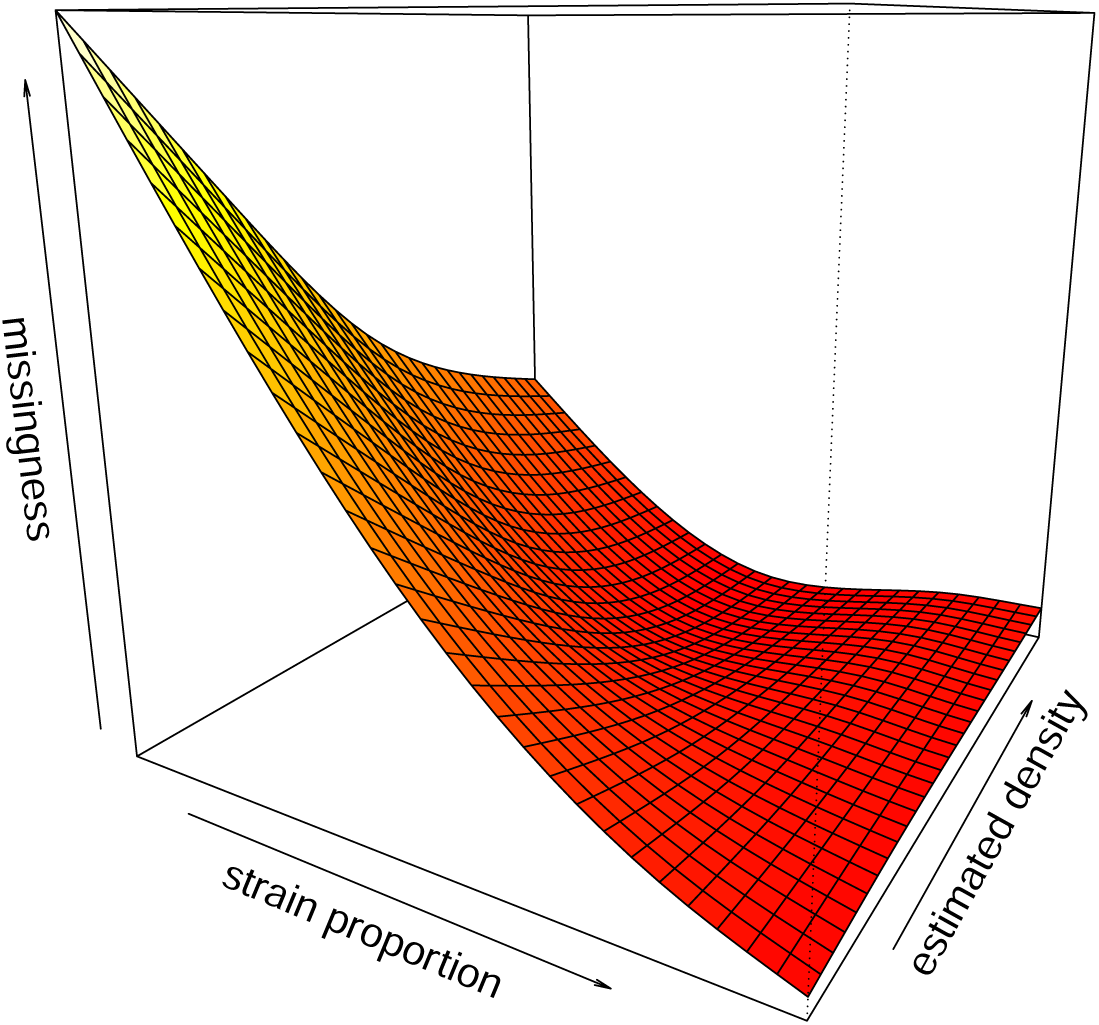
A model fit to predict detection probability from sample parasite density and within-sample strain proportion using empirical data from mixed-strain controls taken from [24]. A general additive model (GAM) with full tensor product smooth was used for the fit, with parasite density and proportion on the log scale. Smaller within-host proportion and overall parasite density resulted in more undetected alleles.

**Figure S.7:**
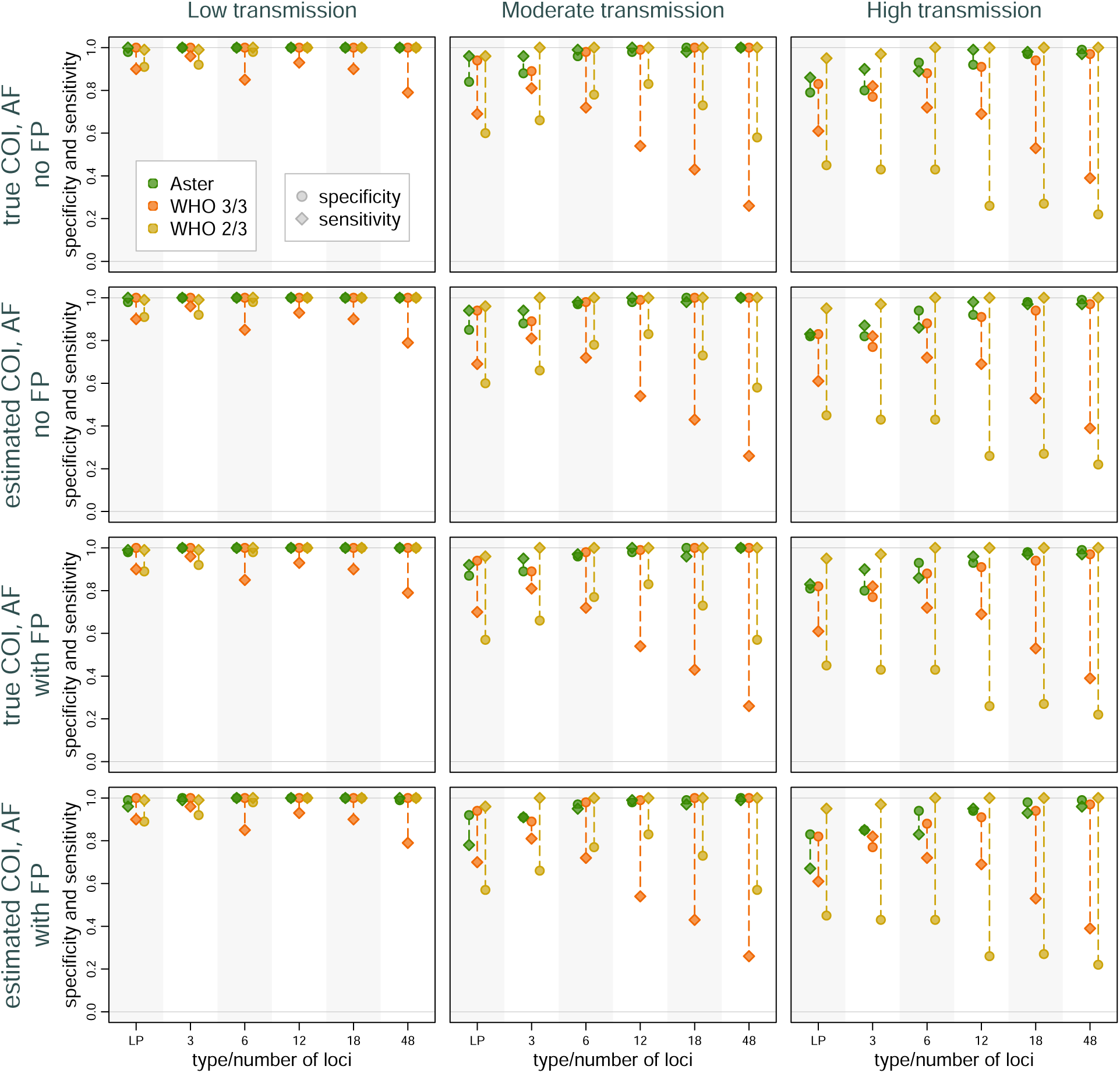
Sensitivity and specificity of Aster and match-counting algorithms exploring the effect of using estimated vs true COI and population allele frequencies (AF) and of adding simulated false positive alleles (FP). Allele frequencies were estimated from 100 D0 samples. False positive rates were 0.1 for the length polymorphism (LP) panel and 0.02 for the amplicon sequencing panels.

**Figure S.8:**
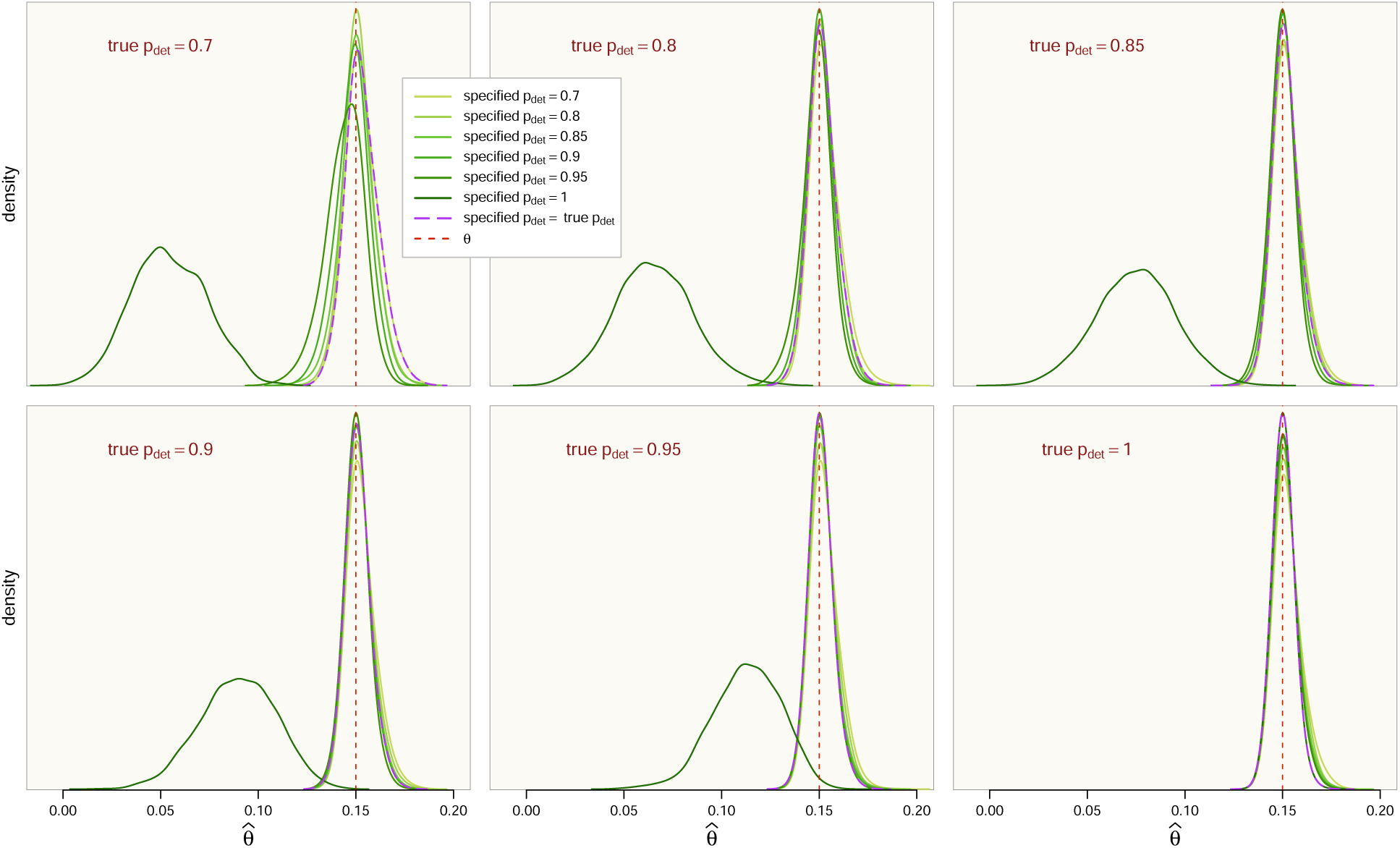
Distribution of failure rate estimates with Aster when using misspecified allele detection probability (*p_det_*). Each panel features one value of true detection probability evaluated with a range of specified values, one of which matched the true probability (purple dotted line marking the corresponding density). Study-level simulation settings were moderate transmission intensity, failure rate (*θ*) of 0.15 (vertical red line), and 0 background relatedness. A 12-locus MAD^4^HatTeR panel was used for the simulations.

